# Differential Regulation of APOE4-Mediated Astrocytic Lipid Metabolism by BMP Signaling Exacerbates Alzheimer’s Disease Pathologies in Neurons

**DOI:** 10.64898/2025.12.14.694212

**Authors:** Anne K. Linden, Amira Affaneh, Aimee Aylward, Yvette Wong, John A. Kessler, Chian-Yu Peng

## Abstract

The two greatest risk factors for Alzheimer’s Disease (AD) are aging and Apolipoprotein E4 (APOE4) polymorphism, yet how these factors interact remain unclear. In this study, we investigate how bone morphogenetic protein (BMP) signaling, which increases with age, contributes to APOE4-induced lipid metabolic dysfunctions using induced-pluripotent stem cell (iPSC)-derived astrocytes and cocultured neurons. Surprisingly, BMP signaling differentially altered lipid droplet formation, cholesterol synthesis and breakdown, and fatty acid-oxidation in APOE4 compared to APOE3 astrocytes, and increased secretion of oxidized LDL (oxLDL). Furthermore, neurons cocultured with BMP4-treated APOE4 astrocytes showed altered transcriptomic profiles based on scRNA-seq as well as increased tau phosphorylation (p-tau). oxLDL treatment similarly increased p-tau and reduced neuronal survival. Conversely, lipid uptake inhibition in neurons rescued the BMP4/APOE4 astrocyte-induced neuronal phenotype. These data demonstrate key interactions between APOE4 and aging-associated molecular signaling in AD pathogenesis and establish a causal linkage between astrocytic lipid metabolism and neuronal tau hyperphosphorylation.

## Introduction

Alzheimer’s disease (AD), the most common neurodegenerative disease, manifests as progressive cognitive decline due to neuronal accumulation of hyperphosphorylated tau tangles, glial accumulation of lipid species, extracellular amyloid plaques, and widespread neurodegeneration[1–6]. While specific mutations may confer an inherited early-onset disease diagnosis, most cases are sporadic and have risk contributions related to both genetics and environment. Apolipoprotein E (APOE) polymorphism is the strongest genetic modifier for late-onset AD (LOAD)[7–12]. While APOE3 (E3) is the most prevalent isoform, patients possessing APOE4 (E4) have a significantly increased risk for AD and have accelerated disease progression[9]. APOE lipidated particles traffic lipids and cholesterol to maintain homeostasis throughout the brain[9, 12–14]. APOE is primarily expressed in the brain in astrocytes, although microglia and neurons also express APOE under specific conditions[15–17]. As systemic lipoprotein particles cannot cross the blood-brain barrier, astrocytic APOE is crucial for the proper trafficking of cholesterol and lipids in support of neuronal health. Lipid dyshomeostasis occurs in LOAD brains[11, 18], and there are disruptions in lipid processing in E4 astrocytes[19–22]. Notably, E4 astrocytes accumulate neurotoxic lipid species that alter normal cell physiology[21, 23]. This suggests that the APOE4-dependent risk of AD may be due, at least in part, to changes in astrocyte lipid metabolism. However, individuals with at least one APOE4 allele do not exhibit pathology until later in life, suggesting some component of the aging process exacerbates the E4 lipid metabolism deficits and accelerates LOAD pathogenesis.

The largest risk factor for LOAD is age, with the majority of patients diagnosed after age 65[24, 25]. Numerous changes in the aging brain, such as compromised autophagy, loss of proteostasis, inflammation, and mitochondrial dysfunction, have been implicated in the pathophysiology of AD, but no specific molecular change has been identified that leads to disease onset and progression[26–28]. Recent proteomic studies have identified Transforming Growth Factor beta (TGFβ) signaling family of proteins as aging regulated factors [29–32]. Specifically, we and other have reported age-related increase in the signaling by TGFβ family member bone morphogenetic protein 4 (BMP4) in both mouse and human brain that correlate with cognitive performance[33–35], and expression of BMP ligands is increased in AD brains compared to age-matched healthy patients[36–39]. Further, inhibition of BMP signaling via overexpression of the BMP antagonist noggin in a *MAPT*^P301S^ mouse model rescues neuronal tau pathology and cognitive deficits[40]. While BMP signaling is typically regarded for its involvement in regulation of cell cycle and lineage commitment during development, it also regulates lipid metabolism and fatty acid oxidation[41–47]. This suggests a possible convergence of age-related enhanced BMP signaling and lipid metabolism pathways relevant to AD. Notably, APOE-BMP interactions have been demonstrated in peripheral organ systems including cardiovascular and adipose tissue[48–52], and altered BMP signaling has been implicated in diseases including atherosclerosis that are also modified by the APOE4 genotype. This prompted us to investigate whether similar interactions between APOE4 and age-associated BMP signaling may be involved in the pathophysiology of AD.

## Results

### BMP signaling induces APOE-specific effects on lipid droplet composition in human iPSC-derived astrocytes

Standard rapid astrocyte differentiation from iPSCs protocol includes BMP4 in the maturation media [53, 54]. To investigate whether BMP signaling interacts with APOE in a genotype-specific manner in regulating astrocytic lipid metabolism, we utilized a modified rapid astrocyte induction protocol without BMP4 in the maturation media to generate human induced astrocytes (iA) from iPSC from heterozygous APOE4 (E4) AD patients versus CRISPR-corrected homozygous APOE3 (E3) isogenic controls[54] (See methods, Figure 1A, Figure S1A). After 35 days in vitro (35DIV), We confirmed that iA express astrocytic molecular markers and capable of glutamate uptake, a critical function of astrocytes (Figure S1B, S1C). No differences in astrocyte marker expression or glutamate uptake were observed between E3 and E4 iA. We then asked if 35DIV iA respond to BMP4 treatment via downstream canonical phospho-SMAD1/5/8 signaling. Following 72 hours of BMP4 treatment, SMAD phosphorylation was significantly increased more than 5-fold in both E3 and E4 iA compared to vehicle-treated controls (Figure 1B, Figure 1C), and there were comparable increases in expression of the BMP-target gene, ID3 (Figure S1D). These findings confirmed that our modified astrocyte differentiation protocol generated BMP responsive iA.

**Figure 1.**
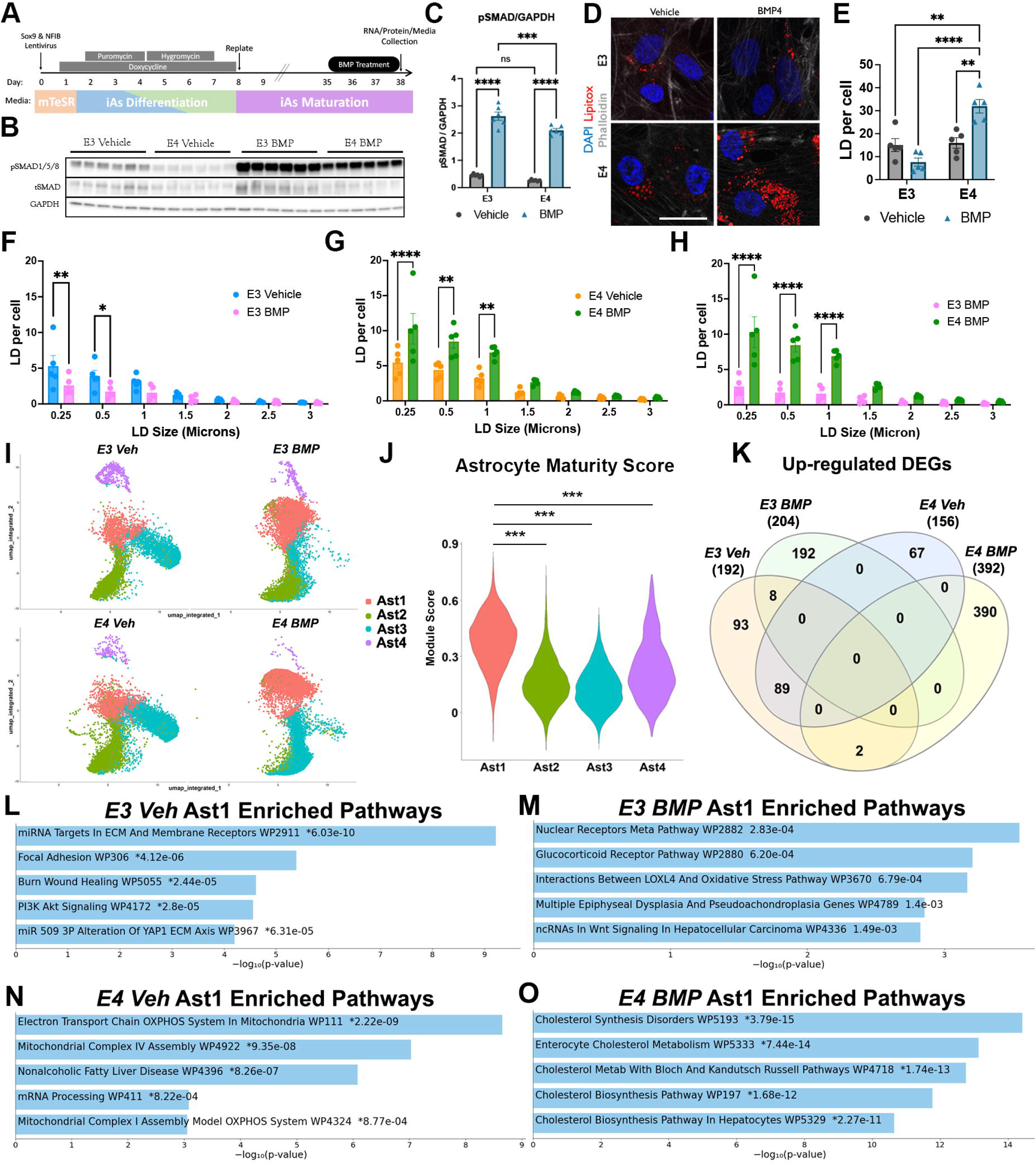
Enhanced BMP-signaling changes in astrocytes are modified by APOE genotype. A) Timeline of iA differentiation and treatment. B) Western blot of phospho-SMAD 1/5/8 and total SMAD in iA groups showing enhanced phospho-SMAD1/5/8 activation upon BMP treatment across both genotypes. C) Quantification of immunoblot in 1B showing enhanced activation upon BMP treatment (n=5). D) Representative image of astrocytes stained for lipid droplets (Blue: DAPI; Red: LipidTOX; White: Phalloidin) scale bar = 25μm. E) Quantification of lipid droplets per cell from 1D (n=5); E4 shows significantly higher lipid droplets per cell. F-H) Quantification of lipid droplets from 1E, showing the number of lipid droplets per cell based on the size of droplets across groups. (n=5). F) Lipid droplets in E3 vehicle and E3 BMP-treated iA. G) Lipid droplets in E4 Vehicle and E4 BMP-treated iA. H) Lipid droplets in E3 vs E4 BMP-treated iA from 1F & 1G. I) UMAP of iA single-cell RNA sequencing across cultures with 4 subclusters. J) Astrocyte maturity score of subclusters based on relative expression of 30 established mature astrocyte markers (gene list found in table S1). K) Venn diagram of uniquely upregulated genes within each culture condition found in scRNAseq. L-O) Enriched pathways based on unique DEGs of Ast1 cluster from scRNA sequencing. Enriched pathways in vehicle-treated E3 iA (L), BMP-treated E3 iA (M), vehicle-treated E4 iA (N) or BMP-treated E4 iA (O) based on unique upregulated DEGs from Venn diagram in K. Data are expressed as mean ± SEM using two-way ANOVA followed by Tukey post-hoc test for multiple comparisons to calculate significance (*p<0.05, **p<0.01, ***p<0.001, ****p<0.0001).

We next examined the effects of BMP signaling on lipid droplet (LD) accumulation in E3 and E4 iA. Using the fluorescent neutral lipid stain LipidTOX to label LDs, we observed significantly greater numbers of LD per cell in E4 iA when compared to E3 iA after 72hrs of BMP4 treatment (Figure 1D, 1E; E3Veh: 15.03; E3BMP: 7.61; E4Veh: 15.9, E4BMP: 31.97). Comparisons of vehicle and BMP4 treated E3 iA showed a reduction of LD numbers with BMP4 treatment, while BMP4 treated E4 iA showed increase LDs when compared to vehicle treated E4 iA. Importantly, contrary to published reports that observed increased number of LDs in E4 iA relative to E3 iA, we did not observe a significant difference in LD numbers between vehicle treated E3 and E4 iA, suggesting that the greater number of LD in E4 iA observed in prior studies is likely is due to the presence of BMP4 in the basal astrocyte maturation media. Lipid droplet size was also influenced by BMP signaling (Figure 1F-H, Figure S1D). Specifically, BMP4 treatment of E3 iA reduced lipid droplet numbers with significantly fewer lipid droplets that were under 1μm (Figure 1F). Conversely, E4BMP iA had significantly greater quantities of lipid droplets in all bins smaller than 1.5μm (Figure 1G, 1H). These findings suggest that BMP signaling induces APOE genotype-specific changes in lipid droplet metabolism, resulting in reduced lipid droplet storage in E3 iA but increased lipid droplet storage in E4 iA.

To understand the molecular pathways contributing to the BMP-induced lipid droplet phenotypes in human astrocytes with distinct APOE genotypes, we performed single-cell RNA sequencing (scRNAseq) of E3 versus E4 iA after 72hr exposure to vehicle or BMP4 (Figure 1I). Data from the four iA experimental groups were normalized and integrated, then analyzed as 4 molecularly distinct cell clusters (Ast1-4) that express known molecular markers of astrocytes (Figure 1J, Figure S2A). Differential expression analysis among Ast1-4 clusters showed that the Ast2 and Ast3 clusters are enriched for markers of glial (NFIA) and neuronal (DCX) precursors (Figure S2A), whereas cells in the Ast4 cluster show high expression of A2-reactive astrocyte markers (Figure S2B)[55]. In comparison, the Ast1 cluster contains cells with the highest expression of 30 established mature astrocyte markers (Figure 1J, Table S1 for gene list). All differential expression and pathway analyses were also performed with cells from each experimental condition considered as one population (pseudobulk) to better align the transcriptomic findings with protein and lipid measurements. In majority of cases, results from Ast1 aligned well with that observed with the pseudobulk expression profiling (data not shown). Therefore, to better understand how BMP signaling affects mature astrocytes in an aging context, we prioritized Ast1 for further analyses.

We identified multiple differentially expressed genes (DEGs) across the 4 experimental groups, followed by pathway analyses using Enrichr[56] and Gene Set Enrichment Analysis (GSEA) with the WikiPathway 2024 human genesets [57, 58]. In comparison to the other 3 experimental groups, the E4 BMP iA group includes at least twice as many unique upregulated DEGs and a similar number of unique downregulated DEGs (Figure 1K, Figure S2B, Table S7), suggesting significant interactions between the two experimental variables. Pathway analyses performed with shared and unique DEGs from the 4 experimental groups revealed that, regardless of APOE genotype, BMP4 treatment reduced expression of genes related to cytoplasmic ribosomal proteins, which is an adaptive transcriptional response that is commonly associated with cellular senescence and stress (Figure S2C, S2D)[59, 60]. Intriguingly, BMP4-treated E4 iA showed a strong enrichment in genes related to cholesterol biosynthesis and metabolism (Figure 1O), which are not present in the other three experimental groups using either 4 group comparisons (Figure 1L-O) or pairwise GSEA analyses (Figure S2C-E). Taken together, the observed changes in lipid droplet size/number and cholesterol biosynthesis gene expression in BMP-treated E4 astrocytes support the hypothesis that BMP signaling differentially regulates cholesterol metabolism in an APOE genotype-specific manner.

As the contents of lipid droplets primarily consist of triglycerides and cholesterol esters, we next compared both relative and normalized expression of all known cholesterol biosynthesis and transport genes, as well as triglyceride and fatty acid biosynthesis genes, across the 4 experimental groups. Strikingly, we found higher relative (Figure 2A) and normalized (Figure 2B) expression of all cholesterol biosynthesis genes in the E4BMP iA when compared to the other 3 experimental groups. By contrast, triglyceride biosynthesis-associated transcripts were higher in both BMP-treated groups compared to vehicle-treated groups (Figure 2C) but were not notably different in the same treatment groups across APOE genotypes when normalized to expression-adjusted controls (Figure 2D). Taken together, these data suggest that BMP signaling preferentially induces increased cholesterol biosynthesis pathways in E4 astrocytes.

**Figure 2.**
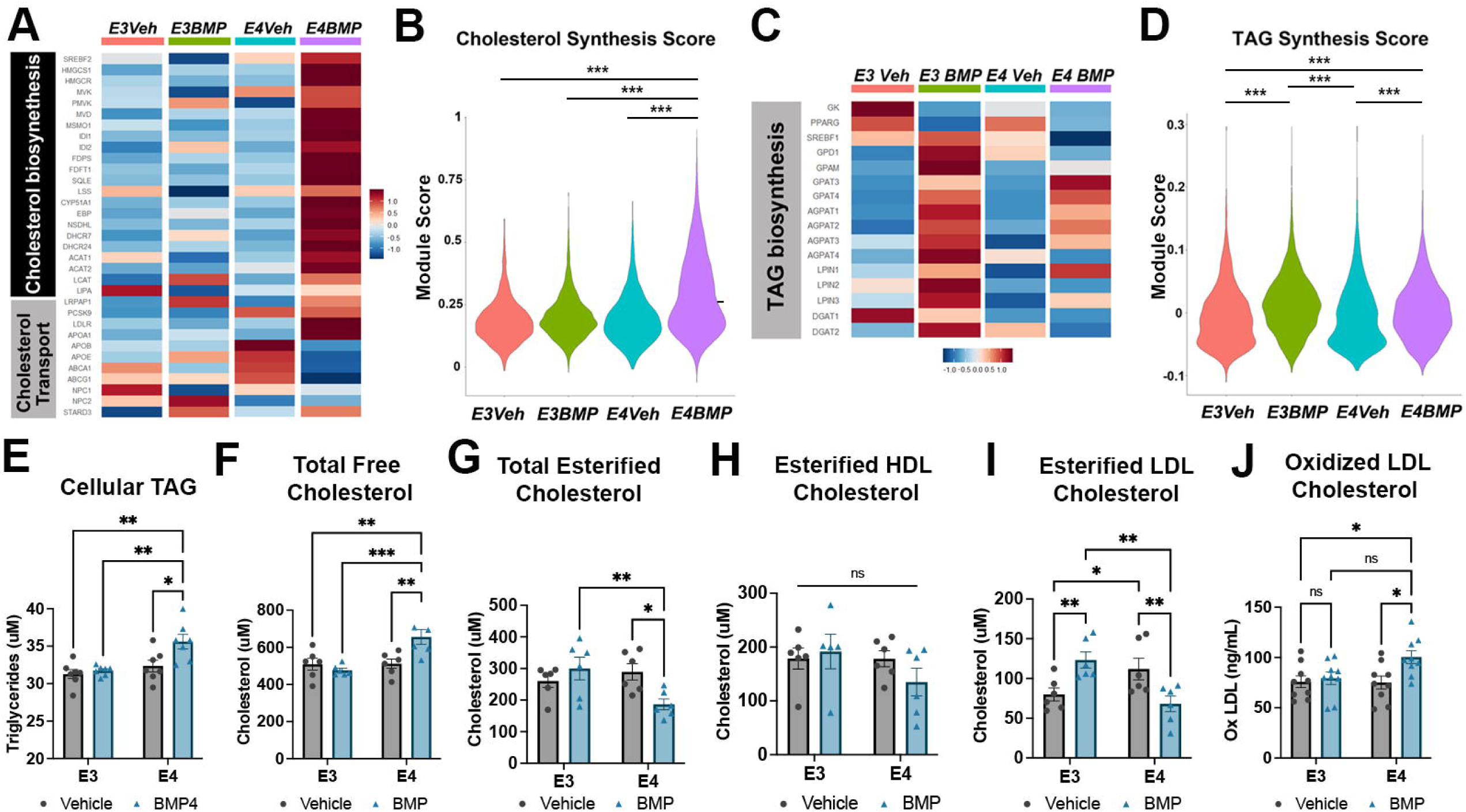
BMP alters lipid and cholesterol composition in APOE4 astrocytes. A) Heatmap of relative expression of cholesterol-related synthesis and transport genes in all iA conditions. B) Violin plot of DEG contributing to enriched cholesterol synthesis in BMP-treated iA from 2A. C) Heatmap of relative gene expression of fatty acid and triglyceride synthesis genes across iA cultures based on scRNAseq. D) Quantification of intracellular triglyceride levels in iA in uM based on Triglyceride-Glo assay. E) Quantification of total free intracellular cholesterol in iA (n=7). F-I) Quantification of intracellular esterified cholesterol in iA (n=6-8); F: total free cholesterol; G: total esterified cholesterol; H: HDL esterified cholesterol; I: LDL esterified cholesterol. J) Quantification of intracellular levels of oxidized LDL in iA (n=9). Data are expressed as mean ± SEM using two-way ANOVA followed by Tukey post-hoc test for multiple comparisons to calculate significance (*p<0.05, **p<0.01, ***p<0.001, ****p<0.0001).

### BMP signaling alters cholesterol composition in APOE4 astrocytes

To determine whether transcriptomic level changes in lipid biosynthesis influence astrocyte lipid homeostasis, we quantified cellular lipids using fluorogenic assays in the 4 experimental conditions. We found that E4BMP iA had significantly higher triglyceride levels (35.61±0.97μM) compared to both E4Veh iA (32.36±0.78μM, p=0.0152) as well as E3 iA regardless of treatment (Figure 2E; E3Veh: 31.38±0.60μM, p = 0.0011; E3BMP: 31.74±0.24μM, p=0.0035). Additionally, fatty acid synthase, FASN, was significantly increased in E4BMP iA (Figure S3B), suggesting upregulation of fatty acid synthesis. Quantification of cellular cholesterol revealed an increase in E4 iA (1244±80.75μM) compared to E3 iA (1109±43.45μM, p=0.0224), although BMP treatment did not further alter the total cellular cholesterol concentration in either genotype (Figure S3A, blue bars. E3BMP: 1120±62.27μM; E4BMP: 1230±43.51μM). While HDL cholesterol comprised the majority of cholesterol content in the iA (Figure S2A, red bars), no difference was observed across experimental groups. In contrast, LDL/vLDL cholesterol levels are higher in E4 iA (426.3±34.98μM) relative to E3 iA (291.4±23.06μM, p=0.0218; Figure S3A), but no significant difference was detected between BMP and vehicle-treated iA with the same APOE genotype (Figure S3A).

To further our understanding of changes in cholesterol metabolism that result from BMP treatment, we directly assessed changes in cellular cholesterol species. We found a significantly higher concentration of free cholesterol in BMP-treated E4 iA (Figure 2F; E4BMP: 655.9±40.25μM; E3Veh: 509.1±31.69μM, p=0.0019; E3BMP: 476.3±11.14μM, p=0.0003; E4Veh: 511.9± 25.22μM, p=0.0022) but significantly lower levels of esterified cholesterol in E4 BMP-treated iA (Figure 2G; E4BMP: 186.4±17.06μM; E4Veh: 289.6±26.0μM, p=0.0104; E3BMP: 299.7±35.91μM, E3Veh: 260.3±20.09μM, p=0.0056). To further assess cholesterol metabolism, we quantified the proportion of HDL vs LDL esterified cholesterol. Most esterified cholesterol was HDL and was not different between iA genotypes in vehicle-treated cells (Figure 2H). However, there were changes in esterified LDL in E3 compared to E4 iA. At baseline, E4 iA have significantly higher levels (111.8±13.56μM) of esterified LDL cholesterol than E3 iA (79.9±8.19μM, p=0.0460) which parallels the increase in LDL shown between E3 and E4 iA (Figure 2I, Figure S3A). BMP treatment significantly increased levels of esterified LDL in E3 iA (123.3±10.11μM, p=0.0091) but conversely decreased levels in E4 iA (68.2±9,83μM, p=0.0086, Figure 2I). These data suggest that in response to BMP treatment, E4 iAs increase free cholesterol and reduce LDL cholesterol esterification, whereas E3 iA increase LDL cholesterol esterification without changing free cholesterol concentrations.

Oxidation of cholesterol can be induced via interaction with free radicals inside the cell and increased LDL species in E4 iA might be at risk of higher oxidation due to free radicals following BMP treatment. We therefore assessed levels of oxidized LDL particles in iAs and found that OxLDL concentrations were significantly increased in E4 iA after BMP treatment (Figure 2J, E4BMP: 100.4±6.34ng/ml; E3Veh: 75.85±5.83ng/ml, p=0.040; E4Veh: 75.12±6.39 ng/ml, p=0.033). To determine whether altered protein expression of enzymes regulating lipid biosynthesis contributes to the observed lipid level changes, we performed western blot analyses of the lipid biosynthesis-associated proteins SREBP2, FASN, DGAT2, and LCAT in the 4 experimental groups. For all proteins examined, we observed an APOE4 genotype-dependent, but not BMP4-dependent, increase in expression (Figure S3C-E), suggesting that transcript level increases observed in E4 BMP iA may be compensatory responses to impaired lipid homeostasis. These data further suggest that the elevated level of cellular lipids in E4 astrocytes may reflect either increased lipid uptake or reduced lipid degradation.

### BMP4 treatment alters lipophagic efficiency in APOE4 astrocytes

Since lysosome-mediated cholesterol degradation can function as a signal to elicit cholesterol biosynthesis, we asked if the increase in LD accumulation and altered cholesterol composition in BMP-treated E4 iA is due, in part, to a change in lipophagic efficiency. We therefore conducted immunocytochemistry on iA following BMP treatment to observe the relation of lysosomes to lipid droplets (Figure 3A). Using LAMP1 to label lysosomes and LipidTOX to stain for lipid droplets, we found that E4 iA contain higher levels (roughly 2-fold) of LAMP1-associated lysosomes than E3 iA (Figure 3B). The overall level of lysosomes did not change upon BMP treatment for either genotype, but the degree of colocalization of lysosomes and LD increased significantly in BMP-treated astrocytes (Figure 3C). BMP treatment significantly increased E3 lysosomal colocalization (Vehicle: 2.32±0.5%; BMP: 14.34±2.54%, p=0.0039). E4 iA had significantly increased LAMP1-LD colocalization at baseline compared to E3 cocultures (E4Veh: 18.36±2.88%, p<0.0001) that was further increased by BMP treatment (E4BMP: 27.21±2.52%, p=0.0305, vs. E3Veh p<0.0001; vs. E3BMP p=0.0014). This suggests that the overall increase of LDs within E4 iA leads to a higher degree of LD targeted for degradation. However, lysosomal acid lipase (LAL), which is necessary for lipid breakdown in lysosomes through lipophagy, was significantly reduced in E4 compared to E3 iA but not altered with BMP4 treatment (Figure 3D). In addition, expression of adipose triglyceride lipase (ATGL), an enzyme that initiates cytosolic lipolysis, was not different among the 4 groups of iA (Figure 3E). This suggests that the relevant degradation pathway that would regulate the accumulation of lipid droplets in E4BMP iA is mediated through lysosomes.

**Figure 3.**
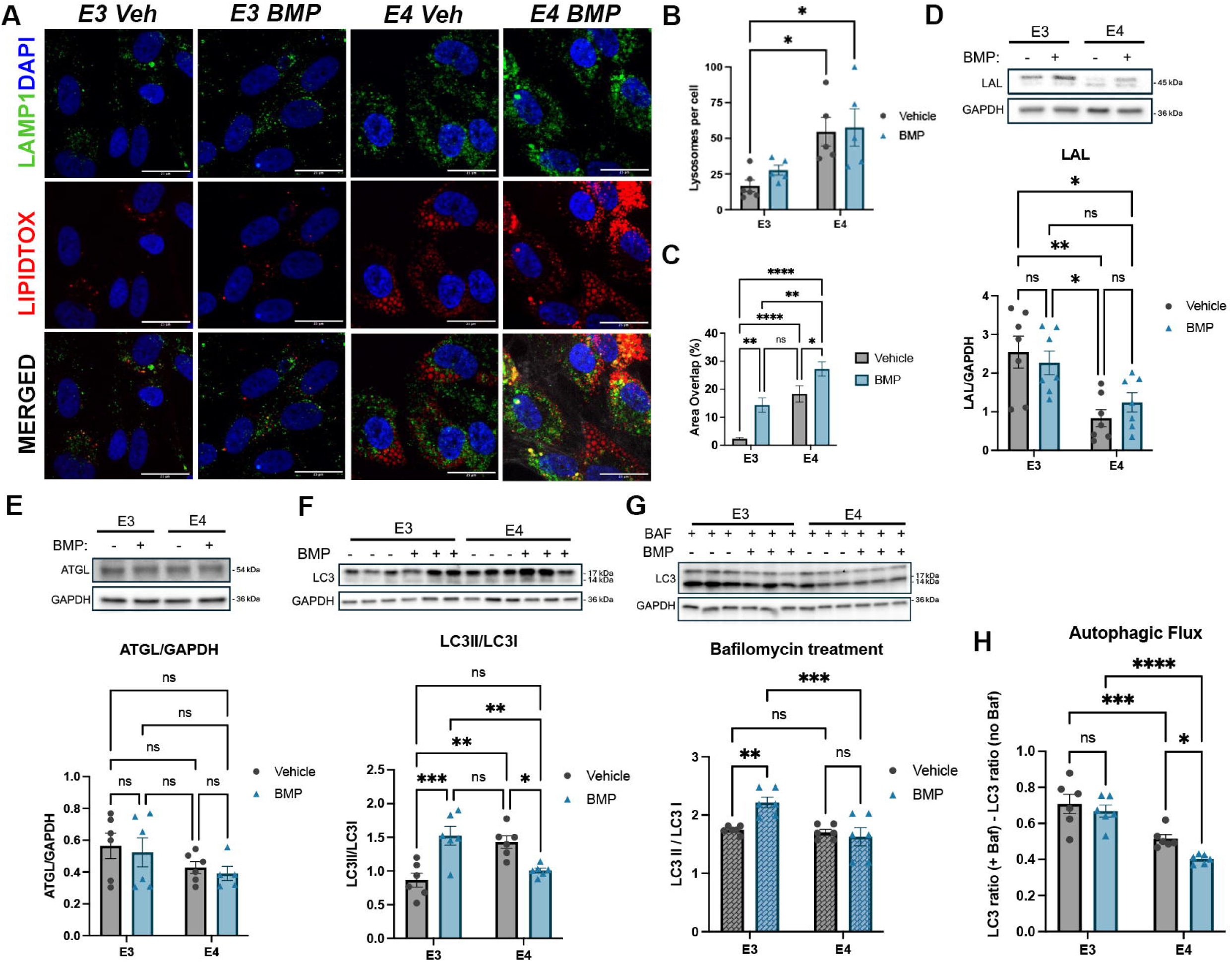
BMP treatment reduces autophagy in APOE4 astrocytes. A) Representative images of immunocytochemistry of iA *Blue: DAPI; Green: LAMP1; Red: LipidTOX;* scale bar= 25μm. B, C) Quantification of immunocytochemistry images (n=5). B) Quantification of lysosomes per cell. C) Quantification of percent overlap between LipidTOX and LAMP1 staining in iA. D) Representative western blot and quantification for LAL (n=6). E) Representative western blot and quantification of ATGL (n=6). F) Representative western blot and quantification of LC3II/LC3I ratio (n=6). G) Western blot and quantification of LC3II/LC3I ratio in iA following 2-hour bafilomycin-A treatment (n=6). H) Autophagic flux calculation based on LC3 ratio with and without bafilomycin treatment (n=6). Data are expressed as mean ± SEM using two-way ANOVA followed by Tukey post-hoc test for multiple comparisons to calculate significance (*p<0.05, **p<0.01, ***p<0.001, ****p<0.0001).

We then asked if the accumulation of LD in E4BMP iA is due to specific downregulation of lipophagic capacity or due to a generalized reduction in autophagy. LC3 conversion from I to II is indicative of the progression of the cell through the autophagy pathway, as LC3II is the more mature form and indicative of autophagosome formation and maturation[61, 62]. At baseline, E4 iA express higher levels of LC3II/LC3I ratio compared to E3 iA (Figure 3F). Following BMP4 treatment, the LC3 II to LC3I ratio increased in E3 iA but reduced in E4 iA. This suggests a reduction in overall autophagic maturation, not just lipophagic efficiency. To test the cell’s capacity for autophagy, we used Bafilomycin A, an inhibitor of autophagy that blocks the fusion of lysosomes and autophagosomes, to inhibit degradation. Following two hours of bafilomycin treatment, the turnover ratio of LC3 was significantly higher in E3BMP iA compared to E3Veh or E4Veh iA (Figure 3G). E4BMP iA did not exhibit a change in the LC3II/LC3I ratio, suggesting that the turnover of autophagosomes does not change in response to LD accumulation. Overall, E3 iA had significantly higher autophagic flux than E4 iA, and the ratio in E3 iA did not change in response to BMP4 treatment (Figure 3H). E4 iA had reduced autophagic flux at baseline, and BMP4 treatment further reduced autophagic capacity..

### Oxidative phosphorylation and fatty acid oxidation in astrocytes are altered by BMP signaling

Astrocytes can utilize lipid components found in LD for energy production via fatty acid oxidation (FAO), a process reported to be reduced in APOE4 astrocytes[20, 22, 63, 64]. Earlier reports have also indicated that BMP signaling promotes FAO in adipose tissue[47, 65]. To examine the effects of BMP signaling on the astrocytic OxPhos and FAO, we performed seahorse analysis on BMP4 or vehicle-treated E3 and E4 iA (Figure 4A). We found that OxPhos in E3BMP iA, as assessed by the oxygen consumption rate (OCR), was significantly elevated (32.28±1.43pmol/min) in comparison to E3Veh (24.21±1.78pmol/min, p=0.036), E4Veh (19.58±1.52pmol/min, p=0.0003), or E4BMP iA (23.88±2.70pmol/min, p=0.0219) (Figure 4A, 4B). No significant differences in basal respiration OCR were detected between BMP4 or vehicle-treated E4 iA (p = 0.398). Similarly, extracellular acidification rate (ECAR) did not differ between vehicle-treated E3 and E4 iA (Figure S4A, S4B). However, BMP4 treatment induced a significant increase in extracellular acidification in E3 iA but not in E4 iA (Figure S4A, S4B), suggesting either glycolytic flux increase and/or increased mitochondrial CO_₂_-driven acidification. When FCCP was applied to induce maximal respiratory capacity of the cells, BMP treatment increased the OCR in both E3 (Figure 4A, 4C, E3BMP: 55.50±2.64pmol/min, E3Veh: 36.49±2.92pmol/min, p=0.0005) and E4 iA (E4BMP: 37.67±3.78pmol/min, E4Veh: 24.92±2.07pmol/min, p =0.0188). E3BMP iA have a significant increase in estimated ATP production through both glycolysis (E3Veh: 56.36±5.50pmol/min; E3BMP: 109.55±8.36pmol/min, p=0.0004) and mitochondrial OxPhos (Figure 4D, E3Veh: MitoATP 98.72±16.3pmol/min, p=0.0056,), as inferred from ECAR and OCR measurements. However, E4 iA only increase ATP production through OxPhos (Figure 4D, E4Veh: 54.29±8.43pmol/min; E4BMP: 91.55±13.20pmol/min, p=0.0259) and not glycolysis (E4Veh: 46.91±4.26pmol/min; E4 BMP:66.69±7.35pmol/min, p=0.4459).

**Figure 4.**
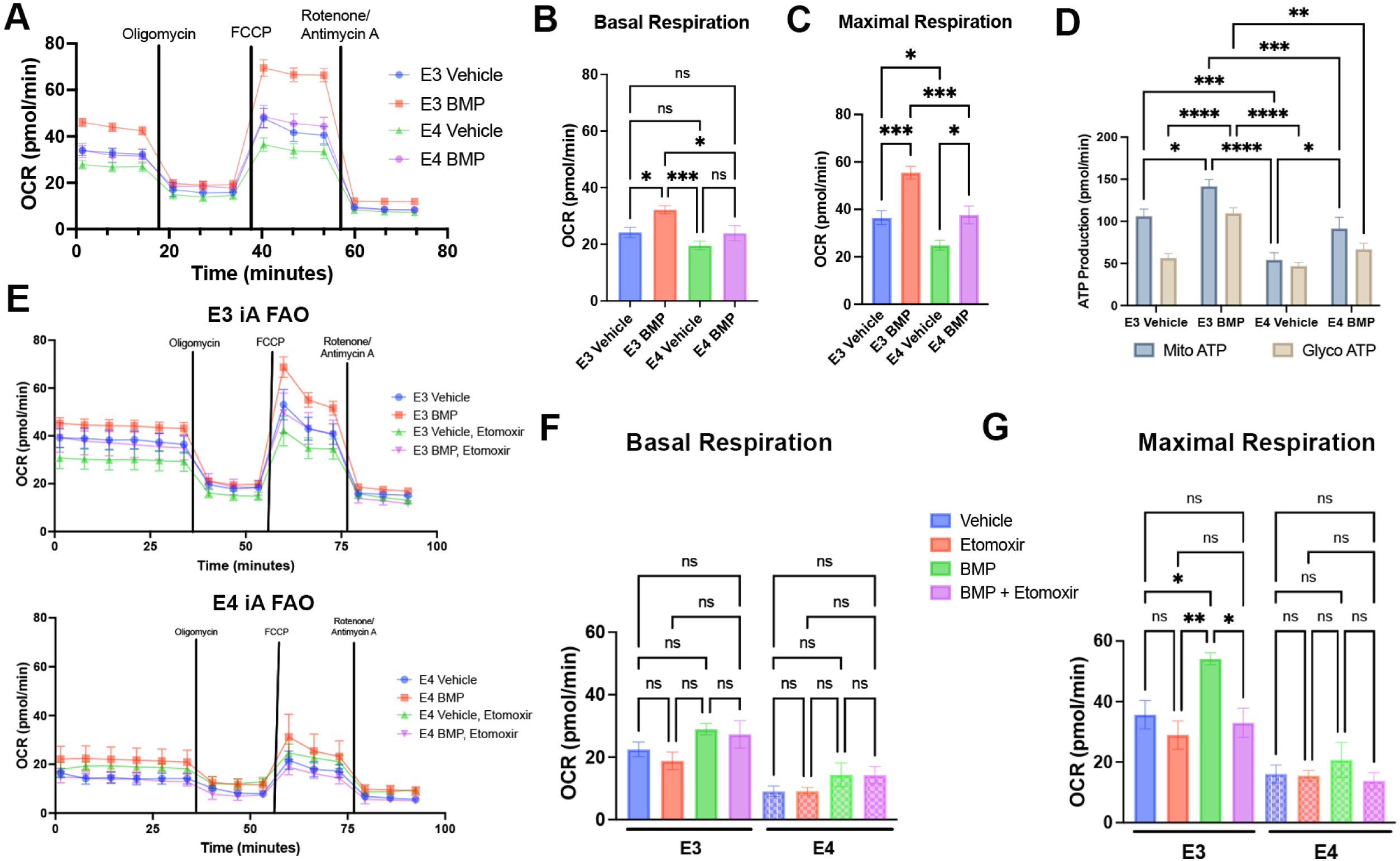
Fatty acid oxidation-dependent enhancement of mitochondrial energy production by BMP signaling is impaired in APOE4 astrocytes. A) Oxygen consumption rate (OCR) of E3 and E4 iA with and without BMP4 treatment during a seahorse-based mitochondrial stress test. Black vertical bars indicate drug treatment during the assay (Oligomycin, FCCP, and Rotenone/Antimycin A (n=8). B) Average basal mitochondrial respiration of iA from the 4 experimental conditions in seahorse analysis before drug treatment in pmol/min. C) Maximal mitochondrial respiration of iAs seahorse analysis following FCCP treatment in pmol/min. D, E) Oxygen consumption rate over time of E3 iA (D) and E4 iA (E) in starved-state conditions with BMP and/or etomoxir treatment (n=6). H) Average baseline reduction of OCR in iA based on etomoxir treatment. F) Average basal mitochondrial respiration of iA under starved-state conditions. G) Average maximal respiration of iA following FCCP treatment in starved-state conditions. Data are expressed as mean ± SEM using two-way ANOVA with repeated measures followed by Tukey post-hoc test for multiple comparisons to calculate significance (*p<0.05, **p<0.005, ***p<0.0005).

We next assessed the FAO contribution to OxPhos capacity of iA under starved conditions with and without Etomoxir, a carnitine transporter inhibitor, to understand the capability for FAO in these cells. Both E3 and E4 iA maintained OxPhos in starved states based on OCR (Figure 4E-4G). BMP4 treatment did not significantly increase basal OCR in starved states in E3 iA (E3Veh: 22.50*±*2.35pmol/min; E3BMP: 28.98±1.82pmol/min, p=0.7295) or E4 iAs(E4Veh: 9.06±1.70pmol/min; E4BMP: 14.37±3.81pmol/min, p=0.9032), suggesting that BMP-mediated increases in basal OCR are glucose-dependent. More interestingly, FCCP treatment in the presence of etomoxir significantly reduced maximal respiration of E3 BMP iA in the starved state (Figure 4G, E3BMP: 54.11±2.00pmol/min; E3BMP + Etomoxir: 32.98±4.82pmol/min, p=0.0112). In contrast, FCCP treatment did not enhance maximal respiration in starved E4BMP iA with or without Etomoxir (Figure 4G, E4BMP: 20.71±5.75pmol/min; E4BMP + Etomoxir: 13.81±2.72pmol/min, p=0.933). These findings suggest that FAO contributes significantly to BMP-induced increases in maximal respiration in starved E3 iA, and that dysfunction in this process impairs energy production in E4 iA under cellular stress. The inability to utilize lipids for mitochondrial energy production, under cellular conditions with enhanced cholesterol biosynthesis, may further explain the lipid accumulation that is observed in E4BMP4 iA.

### BMP signaling-induced lipid secretion by APOE4 astrocytes non-cell autonomously affects neuronal lipid synthesis and metabolism

We next asked whether the changes in lipid homeostatic pathways in E4 iA in response to BMP signaling resulted in altered secretion of cholesterol and lipid components that might influence neighboring cells. We addressed this by assessing changes in astrocyte-conditioned media (ACM) following 72 hours of vehicle or BMP4 treatment. We first examined the levels of a select panel of chemokines and cytokines in ACM and found no significant changes apart from a small increase in IL6 in E4 ACM (Figure S5A-E). Given that APOE can be associated with amyloid species, we also assessed levels of amyloid in iA and ACM following BMP treatment (Figure S5F-S5J, S5C). No change in amyloid β expression was observed in iA regardless of treatment (Figure S5F, S5G). Although there was a slight increase in amyloid β42 species in E4 astrocyte media compared to E3 astrocyte media, we did not find a difference in the amyloid 40-42 ratio among iA groups (Figure S5H-S5J). This suggests that amyloid processing or secretion in astrocytes is not significantly impacted by BMP signaling.

We next assessed changes in secreted lipids between these conditions. Assessment of triglyceride levels in the ACM revealed significant increases in E4BMP (Figure 5A, 33.54±0.962μM) in comparison to E3Veh (28.32±0.579μM, p =0.0006), E3BMP (28.78±1.12μM, p =0.0007), or E4Veh ACM (28.49±0.59μM, p =0.0008). In contrast, total HDL (Figure 5B) and LDL (Figure 5C) levels of ACM cholesterol remained the same across all conditions. Although we detected altered levels of cellular cholesterol ester in E4 BMP iA, no significant differences in cholesterol ester levels were observed in the media among the 4 experimental groups (Figure 5D). Based on increased levels of cellular oxLDL in iA, we next examined if oxLDL secretion is altered. E4Veh ACM had significantly elevated levels of oxLDL compared to E3Veh ACM (Figure 5E, E3Veh: 92.40±7.52 ng/ml, E4Veh: 118.3±5.45 ng/ml, p=0.0385). More importantly, BMP treatment significantly increased oxLDL in both E3 (E3BMP: 117.6±6.13 ng/ml; E3Veh: 92.40±7.52 ng/ml, p=0.0430) and E4 (E4BMP: 142.9±12.64 ng/ml, E4Veh: 118.2±5.45 ng/ml, p=0.0479) ACM. BMP4 treatment of E3 iA increased secreted oxLDL levels roughly to baseline E4 iA levels, whereas BMP4 treatment of E4 iA further increased extracellular oxLDL to levels significantly higher than in any other condition.

**Figure 5.**
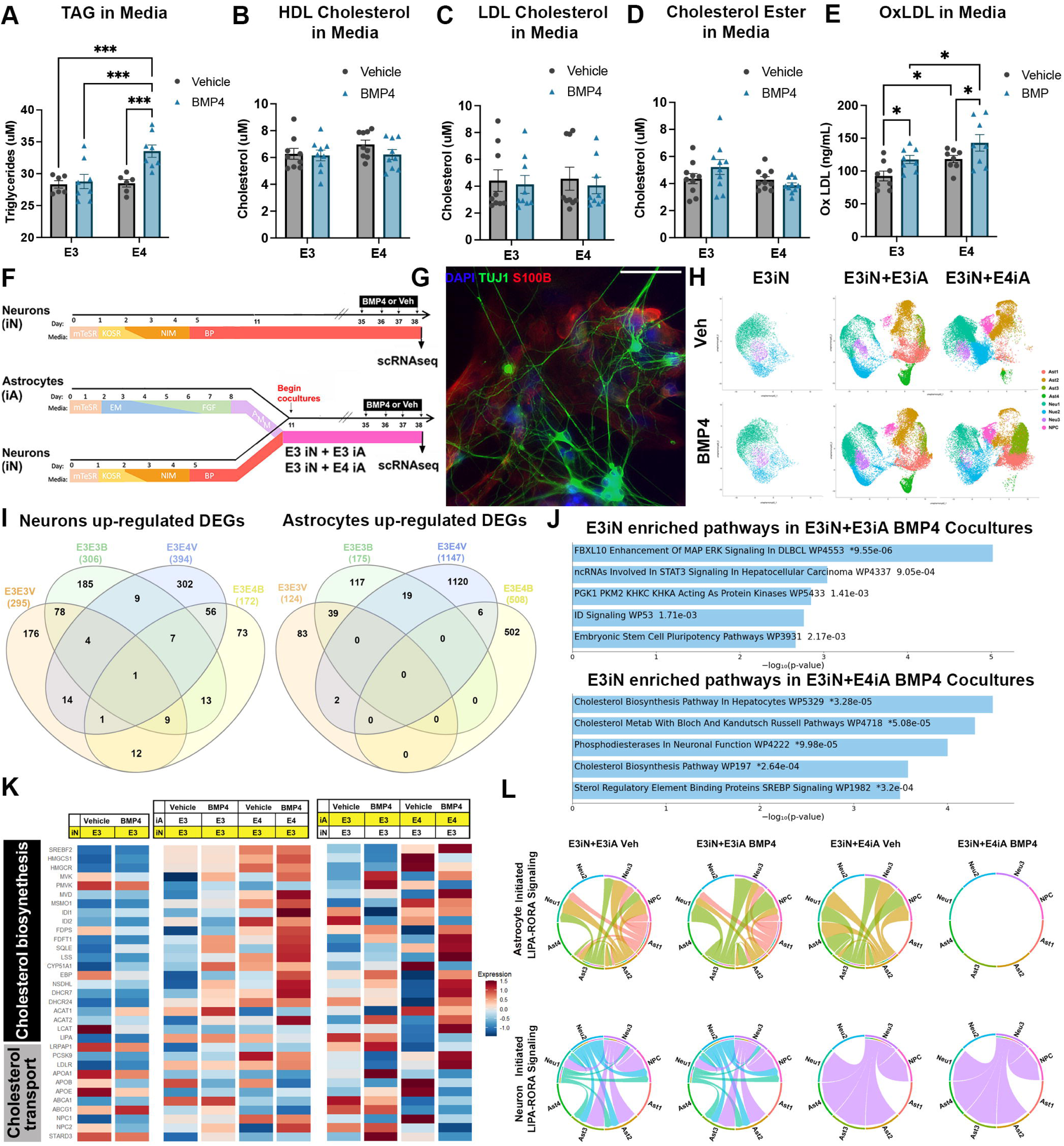
BMP signaling alters APOE4 astrocytic lipid secretion and drives neuronal lipid dysfunction in cocultures. A) Quantification of triglycerides in ACM in *μ*M. B-E) Quantification of secreted cholesterol species in ACM; B: total HDL cholesterol in ACM, C: total LDL cholesterol in ACM, D: total esterified cholesterol in ACM, E: Oxidized LDL cholesterol in ACM in ng/ml. Data of A-E are quantified with n=8-9 per condition and expressed as mean ± SEM using two-way ANOVA followed by Tukey post-hoc test for multiple comparisons to calculate significance (*p<0.05, **p<0.01, ***p<0.001, ****p<0.0001). F) Coculture timeline of iA and iN for sequencing experiments. G) Representative image of cocultures. (Blue: DAPI; Green: TUJ1; Red: S100B) scale bar = 100μm. H) Representative UMAP of scRNAseq data sets from E3iN, E3iN+E3iA, and E3iN+E4iA under vehicle and BMP conditions. I) Venn diagram illustrating number of unique and overlapping up-regulated DEGs in E3 Neu1 cluster neurons and E3 or E4 Ast1 cluster astrocytes from each of the four coculture experimental groups. J) Differentially enriched pathways in E3 Neu1 cluster iN cocultured with either E3 iA or E4 iA in the presence of BMP4. K) Heatmap representation of the relative expression of genes associated with cholesterol biosynthesis and transport in iNs compared across 6 experimental conditions. The same geneset is also compared in iA across the 4 cocultured conditions. L) Chord plot representation of astrocyte (top) and neuron (bottom) initiated cell-cell interactions based on expression of LIPA and RORα transcript levels in each cell cluster of the 4 coculture conditions.

These observations raised the possibility that increased oxLDL secretion by E4 astrocytes in response to BMP signaling might differentially influence neighboring neurons. To address this issue, we first examined the effects on the neuronal phenotype of E3 neurons (E3 iN) cocultured with E4 versus E3 iA. We independently generated E3 iN, E3 iA, or E4 iA and subsequently combined E3 iN with E3 or E4 iA on day 11 post-differentiation as co-cultures, until day 35 when we treated co-cultures with BMP4 or vehicle for 72hrs (Figure 5F). S100B and TUJ1 immunostaining were performed to confirm astrocytic and neuronal identities respectively (Figure 5G).

To examine neuronal transcriptomic level changes, we conducted scRNAseq analysis in E3 iN only cultures, E3iN+E3iA, and E3iN+E4iA cocultures. Cell clustering and identity analyses of the cocultures revealed three neuronal and four astrocytic populations with expression profiles that aligned with astrocytic or neuronal subtypes identified in the single-cell type cultures (Figure 5H). Examination of neuronal subtype marker expression in iN-only cultures (Figure S5B) as well as iN+iA cocultures (Figure S6A) confirmed iN express excitatory glutamate receptor and cortical identity markers (Figure S5B), consistent with the generation of cortical excitatory neurons that are expected from the differentiation protocol.

To identify astrocyte-induced changes in neurons, we first examined baseline effects of BMP4 treatment in E3 iN-only cultures. DEG analysis identified a relatively small number of upregulated genes in E3 BMP iN, with most enriched pathways related to BMP/TGFβ signaling activation (Figure S6B, S6C, Table S2). More importantly, we did not observe any transcript level changes related to cholesterol biosynthesis in E3 BMP iN-only cultures (Figure S6D, 5K). We next investigated whether BMP signaling in astrocytes differentially regulates neuronal gene expression in neuron-astrocyte cocultures. DEG of vehicle and BMP4 treated E3iN+E3iA and E3iN+E4iA cocultures revealed similar numbers of unique DEGs in E3iN+E3iA cocultures with and without BMP4. In contrast, significantly more DEGs in both neurons and astrocytes of the E3iN+E4iA Veh group are reduced by BMP treatment (Figure 5I). Pathway analyses of cocultured iA revealed that, BMP4-treated E4 iA in cocultures show elevated cholesterol biosynthesis transcripts compared to cocultured iA in the other experimental groups (Figure 5K, S7B), albeit to a lesser degree compared to iA-only cultures.. Interestingly, pathway analyses with unique DEGs in the E3 iN from the cocultures revealed an enrichment in cholesterol biosynthesis only in E3 iN cocultured with E4 iA and BMP4, and not in E3 iN in any other coculture condition (Figure 5J, 5K). Together, these findings demonstrate that non-cell autonomous effects of BMP signaling in E4 iA may further regulate neuronal cholesterol biosynthesis.

To help define specific lipid-related cell-cell interactions between iA and iN that contribute to the non-cell autonomous effects, we performed ligand-receptor pair inference analysis using the CellChat algorithm with a focus on cholesterol-mediated signaling. We found that, unlike other experimental groups, E3iN+E4iA BMP cocultures showed a loss of reciprocal interactions mediated by expression of lysosomal acidic lipase transcript LIPA and expression of retinoid orphan receptor alpha (RORα) (Figure 5L). This finding is consistent with the impaired lysosomal lipid degradation phenotype observed in iAs (Figure 3) and suggests that nuclear receptor signaling, which is known to modulate cholesterol efflux through transcriptional regulation of ABCA1[66–68], may play a role in the neuronal transduction of astrocytic-initiated lipid signals in E3iN+E4iA BMP cocultures.

### Astrocytic OxLDL secretion reduces neuronal viability and increases tau phosphorylation

To specifically determine the effects of BMP-induced astrocytic lipid changes on neuronal health, we first tested the resiliency of neurons to oxLDL, a lipid species that is secreted at higher levels by E4 BMP iA (Figure 5E). We used live cell imaging to track neuronal viability over time after exposure to oxLDL. Neurons were treated with oxLDL, or a lipid uptake inhibitor (LUI, scavenger receptor class B type 1 inhibitor) to prevent lipid uptake in neurons, or co-treated with both oxLDL and LUI (Figure 6A). Quantification of live cells/total cells normalized to time 0 revealed significant vulnerability of neurons to cell death when exposed to oxLDL (Figure 6B). Within just 6 hours of oxLDL treatment, neuronal survival was significantly decreased in cultures treated with oxLDL (73±4.11%) compared to vehicle-treated neurons (96.34±0.05%, p<0.0001) or neurons co-treated with oxLDL and LUI (95.64±0.21%, p<0.0001). After 24 hours of oxLDL treatment, neuronal survival was reduced by roughly 30% compared to the other culture groups. Importantly, the effect of oxLDL on neuron death was blunted when co-treated with LUI (Fig. 6B). Thus, neurons are vulnerable to oxLDL exposure, but cell viability can be rescued by inhibiting oxLDL uptake.

**Figure 6.**
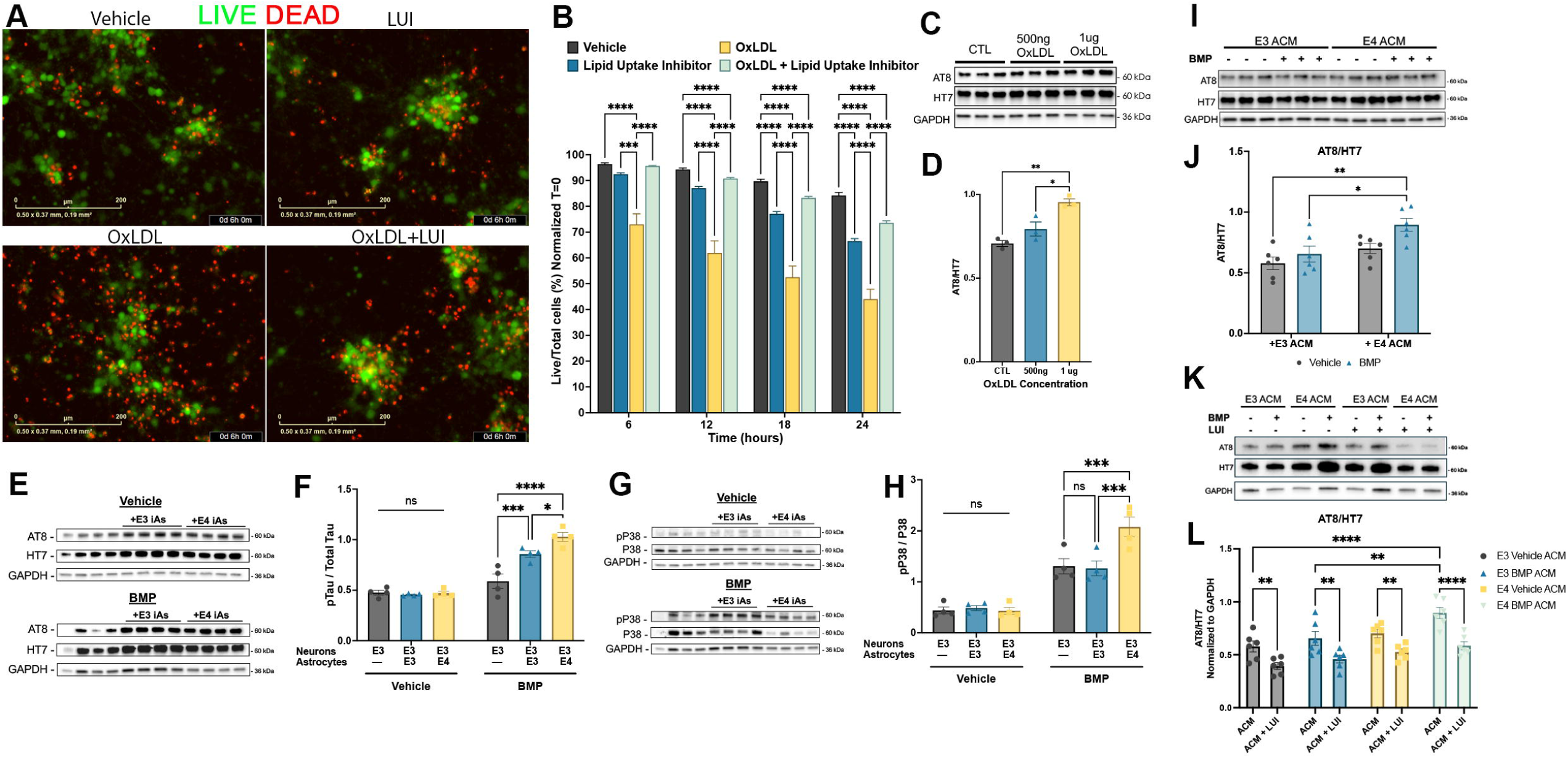
OxLDL released from BMP-treated APOE4 astrocytes increases tau phosphorylation, reduces cell viability, and is recovered with lipid uptake inhibition. A) Representative fluorescent images of Calcien AM (green) and cytotox red (red) as measures of live and dead iN at 6h post-treatment with OxLDL, LUI, or co-treatment. Scale bar = 200 μ*m*. B) Quantification of cell viability of cells at 0, 6, 12, and 24h time points following treatment based on the percent of the live cell over total cells in the frame normalized to time 0h (n=3). C) Representative western blot and quantification of pTau / total Tau protein as labeled by AT8/HT7 antibodies in iN following vehicle, 500ng, or 1μg oxidized LDL treatment for 72 hours (n=3). E) Representative western blot and quantification of pTau / total Tau in iN following coculture with E3 or E4 iA in control or BMP4-treated conditions. F) Quantification of western blots from E (n=4). G) Representative western blot and quantification of pP38/P38 in iN following coculture with E3 or E4 iA in control or BMP4-treated conditions. H) Quantification of western blots from G (n=4). I) Representative western blot and quantification of pTau / total Tau protein in iN following ACM treatment for 72 hours. J) Quantification of western blot from I (n=6). K) Representative western blot of pTau / total Tau protein from iN following ACM treatment alone or with co-treatment of a lipid uptake inhibitor (LUI) for 72 hours. L) Quantification of pTau expression in K (n=6). Data in B are expressed as mean ± SEM using two-way ANOVA with repeated measures, followed by Tukey post-hoc test for multiple comparisons to calculate significance. All other data in the figure are expressed as mean ± SEM using two-way ANOVA followed by Tukey post-hoc test for multiple comparisons to calculate significance (*p<0.05, **p<0.01, ***p<0.001, ****p<0.0001).

To understand potential mechanisms through which oxLDL may affect neuronal viability, we assessed tau phosphorylation in response to oxLDL exposure. Incubating iN with increasing amounts of oxLDL for 72 hours resulted in increased pTau in neurons based on immunoblotting for AT8 expression (Figure 6C, 6D). This suggests that neurons are sensitive to increased levels of oxidized cholesterol species and respond in part by increasing Tau phosphorylation. We then assessed if indirect coculture of iAs and iN without direct cell-cell contact between the two cell types (see Methods) modulates tau phosphorylation in a similar manner. Indirect coculture of iN and iAs did not influence tau phosphorylation at baseline (Figure 6E and 6F). However, BMP treatment of cocultures significantly increased pTau in E3 iN compared to vehicle conditions (Figure 6E and 6F). Moreover, co-culture with E3 iA after BMP treatment further increased pTau levels, which was even further increased in co-cultures with E4 iA (Figure 6E and 6F), demonstrating that E4 astrocyte secretion is sufficient to significantly elevate phosphorylated tau levels in neurons under BMP signaling conditions.

Non-canonical BMP signaling via P38 activation has been shown previously to potentially mediate increases in pTau in E4 iN, but not E3 iN^35^. We therefore asked whether P38 activation may mediate BMP-induced pTau expression in E3 neurons under coculture conditions. We found that BMP treatment influenced P38 expression in E3 iN cultured indirectly with E4 iA with higher levels of phospho-P38 than other cultures treated with BMP (Figure 6G and 6H). These results suggest that BMP treatment of cocultures elicits neuronal tau phosphorylation in part due to increased levels of P38-mediated activation downstream of BMP signaling.

Under coculture conditions, responses to BMP signaling may be due to non-cell autonomous feedback between cell types. To assess the direct role of secreted astrocytic factors on neuronal phenotypes, neurons were incubated in astrocyte conditioned medium (ACM) obtained from either E3 iA or E4 iA for 72hrs to avoid secondary feedback from neuronal responses. Importantly, iN exposed to ACM from E4BMP iA exhibited increased tau phosphorylation based on AT8 immunoblotting (Figure 6I and 6J). This suggests that secreted factors found within ACM from E4 BMP iA could alone elicit tau phosphorylation responses in neurons. we then asked if BMP-induced changes in tau phosphorylation could be inhibited by directly blocking neuronal oxLDL uptake. ACM from either E3 or E4 iA was introduced in the presence or absence of an LUI before neurons were analyzed for tau phosphorylation. Importantly, co-treatment of LUI significantly blunted changes in pTau expression in iN in the presence of ACM (Figure 6K and 6L). In fact, all ACM-treated iN had reduced pTau expression when cotreated with LUI, suggesting that even baseline neuronal lipid uptake influenced tau phosphorylation in neurons.

## Discussion

Studies of LOAD pathogenesis have utilized genetic risk factors to understand changes in cellular physiology that may precede cognitive deficits. However, these studies often fail to include aging components as a trigger for pathological cellular cascades. The current study sought to understand how BMP signaling interact with APOE genotype to alter astrocytic lipid metabolism and resultant neuronal pathophysiology. Our findings suggest that the age-related increase in BMP signaling may be one of the physiologic changes that trigger the onset of LOAD. Alois Alzheimer’s original description of AD described “adipose saccules” (lipid droplets) in glial cells surrounding amyloid plaques as a key neuropathological feature of AD. However, how normal aging contribute to the abnormal lipids to neuronal degeneration has been unclear. The APOE4 polymorphism in humans and in targeted replacement mice leads to increased total cholesterol levels in blood and peripheral tissues, specifically levels of LDL cholesterol lipoparticles[69, 70]. APOE4 astrocytes similarly exhibit increased cellular cholesterol levels despite the blood-brain barrier, suggesting that the increased levels reflect abnormal synthesis, uptake, or processing of cholesterol by astrocytes[71, 72]. Our study found that BMP signaling in APOE4 astrocytes increased lipid droplet formation and lipid homeostasis along with increased cellular cholesterol, secretion of triglycerides and oxLDL. Furthermore, we demonstrated that astrocytic release of lipids, including oxLDL, results in increased tau phosphorylation and reduced neuronal survival, indicating a role for lipid dyshomeostasis in the neurodegenerative process. The increase in cholesterol and fatty acid synthesis gene expression in E4 BMP iA despite high baseline cellular cholesterol levels indicates a potential problem with the sensing of newly synthesized cholesterol or its targeting to the appropriate compartment within the astrocyte for its intended functional purpose.

Previous studies on the role of APOE in intracellular trafficking of lipids have reported impaired targeting of APOE4-associated cellular lipoparticles to the ER as well as lysosomal-autophagic compartments[72, 73]. We found that BMP signaling further reduces autophagic flux in E4 iA, coinciding with impaired lipid droplet breakdown through lipophagy. This is supported by our scRNAseq findings that E4BMP iA have reduced expression of LIPA, which encodes a lysosomal acidic lipase that is necessary for breaking down lipid droplet-associated cholesterol esters. Lysosomal lipid degradation plays a key role in the transcriptional regulation of cholesterol biosynthesis, in part through NPC1/2-mediated transport of lysosomal cholesterol to the ER where the cholesterol sensing-SCAP protein regulates the ER retention or nuclear translocation of the master cholesterol biosynthesis transcription factor SREBP2[74–76]. It is possible that loss of LAL-mediated lipophagy in E4BMP iA led to reduced lysosomal and ER cholesterol levels, which in turn, resulted in SREBP2 nuclear translocation and increased cholesterol synthesis despite higher cellular levels of cholesterol or triglycerides in E4BMP iA. Future studies would benefit from understanding the direct BMP-regulated molecular mechanism by which cholesterol is targeted to specific organelles for processing and metabolism. Elevated cholesterol within cells has also been shown to increase the peroxidation of LDL particles[73], an effect observed in E4BMP iA. The increased susceptibility of LDL to oxidation may partly explain the elevated oxLDL seen in E4 astrocyte cultures at baseline and further exacerbated by enhanced BMP signaling. Notably, our study discovered a direct, novel link between astrocyte-derived oxysterol secretion and phosphorylated tau elevation in neurons. Further, we found that oxLDL reduces survival of iPSC-derived AD neurons. While oxLDL toxicity has previously been suggested to play a role in LOAD[76], our findings specifically demonstrate the neurotoxic effects of astrocyte release of the molecule. Thus, the BMP-mediated increase in oxLDL may serve as a convergence point between aging-associated and APOE-mediated disease pathology in LOAD.

Our investigation into the role of BMP4 in astrocyte mitochondrial function suggested that BMP4 enhances mitochondrial oxidative capacity in E3 astrocytes via increased ATP-linked respiration and spare capacity, in part through substrate flexibility remodel via increased fatty acid oxidation. The observed increase in energy production at the 72hr time point in BMP4 treated E3 iA is consistent with reports of TGFβ family proteins initiating metabolic remodeling by transiently increase energy production in response to cellular stress, differentiation, or senescence signals[77–80]. In contrast, E4 iA were incapable of overcoming energetic demand via fatty acid oxidation. This finding is also consistent with the reduced lysosomal degradation of lipids we observed in E4BMP iA, leading to lower fatty acid utilization for FAO. Notably. oxidative phosphorylation is decreased across age in a variety of human and animal models[27, 81–84]. Since the age-related decrease in mitochondrial respiration efficiency parallels an increase in BMP4 expression in aged animals, future studies that elucidate the casual relationship between sustained BMP signaling activation and astrocytic mitochondrial bioenergetic efficiency may provide insights into aging-associated glial dysfunctions.

To help define the role of BMP-mediated lipid metabolism changes in iA on the neurodegenerative process, we examined non-cell autonomous effects on neurons resulting from astrocytic lipid dyshomeostasis. Our observation that E4BMP iA released increased levels of oxLDL led us to examine the effects on neurons of exposure to exogenous oxLDL, and found that it reduced neuronal viability and increased tau phosphorylation. Effects of oxidative stress, including lipid peroxidation, on neuronal viability and tau phosphorylation has been previously reported[85–87]. In fact, oxLDL signaling through the CD36 scavenger class B receptor has been shown to elicit MAPK signaling pathways that can directly contribute to tau phosphorylation[88]. This suggests a specific pathway through which BMP-mediated enhanced oxLDL concentrations in E4 ACM may directly influence tau pathology in cocultured neurons. ACM from E4 BMP iA increased tau phosphorylation in neurons, and this increase was inhibited by co-treatment with an LUI. This further supports the notion that astrocyte-secreted lipid species, including oxLDL, contribute to BMP-induced neuronal tau pathology. Interestingly, indirect coculture of neurons with E4 astrocytes with BMP treatment elicited phosphorylation of p38 MAPK, a kinase that can directly phosphorylate tau, suggesting a possible secondary mechanism through which BMP-treated E4 iA may induce tau phosphorylation metabolism changes that occur in LOAD pathology. in neurons.

Our study does possess some limitations that may impact the overall translatability of results. Firstly, the lentiviral-induced protocol for *in vitro* generation of astrocytes and neurons, while consistent in generating specific cell types, likely does not fully recapitulate the mature phenotype of cells in an aged brain. Further, while we utilized a 72-hour protocol for enhanced BMP treatment, that does not fully recapitulate the progressive upregulation of BMP signaling that is seen across age over time. There are likely compensatory cellular changes that occur as a result of chronic enhanced BMP signaling across age that are not reproduced in the current model. Lastly, while E3 neurons saw a negligible change in phosphorylated tau in response to BMP treatment in our previous study[40], we cannot rule out the possibility that ACM treatment or coculture conditions that included BMP4 did not elicit neuronal-specific responses independent of astrocytic phenotypes. Future studies could utilize cell-specific constitutively active BMP receptors in astrocytes or BMP receptor knockout viruses in neurons to dissect cell autonomous versus non-cell autonomous effects of BMP signaling activation both in cocultures and transgenic mouse models of LOAD. Collectively, our findings provide additional insights into how the age-associated increase in BMP signaling and APOE genotypes interact to regulate astrocytic lipid metabolism and how these changes contribute to the pathophysiology of LOAD.

## Acknowledgments

This work was supported by funding from the Alzheimer’s Association’s AD Strategic Funds (ABA-22-974612) and NUseq 2022 and 2023 Illumina pilot grants awarded to C.-Y.P. A.K.L was funded by the NIH-NIA training grant F31AG079540. The viruses were produced by the Northwestern University Skin Biology and Diseases Resource-based Center (P30AR075049) with support from NIH/NIAMS. RNA sequencing by the NUseq Core Facility were supported by funding from the NIH Grant 1S10OD025120. Seahorse data were obtained in the Metabolomics Core Facility at Robert H. Lurie Comprehensive Cancer Center of Northwestern University. IncuCyte Live-cell analysis data collection was conducted thanks to the ANTEC Core Facility of the Center for Regenerative Nanomedicine at Northwestern University. We would also like to thank Anil Wadhwani for his work in isogenic CRISPR/Cas9 correction of patient cell lines and generous ongoing support from Mark and Mary Nagelvoort.

## Author Contributions

A.K.L., J.A.K., and C.-Y.P. contributed to the conception and design of the study. A.K.L., A.A. A.J.A., and C.-Y.P. contributed to the acquisition and analysis of data; A.K.L., Y.W., J.A.K., and C.-Y.P. contributed to experimental design and interpretation of the data. A.K.L., A.A., J.A.K., and C.-Y.P. contributed to drafting the text or preparing the figures.

## STAR Methods

### iPSC culture

Induced pluripotent stem cell (iPSC) lines were generated from fibroblasts of three Alzheimer’s Disease patients with APOE3/E4 genotype and (AD1: AG11414, AD2: AG05810, and AD3: AG04402, from Coriell Institute for Medical Research). Fibroblasts were converted into induced pluripotent stem cells as described previously[89], and isogenic controls were generated via CRISPR-based correction to generate APOE3/3 control lines (AD1: G10, AD2: D6, AD3: H7) as previously described[17, 40]. IPSCs were maintained with MteSR1 media (Stemcell Technologies, 100-0276) on Matrigel-coated plates (Corning, 353046) and passaged routinely at ∼80% confluency with ReLeSR (Stemcell Technologies, 100-0483) for stem cell maintenance (Stemcell Technologies, 05872). IPSC cultures were maintained in a 37C incubator with 5% CO2.

### Viral transduction

IPSC cultures were cleaned of spontaneous differentiation before single cell dissociation with accutase (Stemcell technologies, 07920) for 10 minutes followed by 1:1 quenching with mTeSR. Cells were pelleted at 800 RPM for 5 minutes before resuspension with mTeSR and 10 uM Rock inhibitor, Y-27632 (Reprocell, 04-0012-02), and lentiviruses encoding rtTA as well as pTetO-Ngn2-puro for neuronal cultures or pTetO-Sox9-puro and pTetO-Nfib-hygro for astrocytes (plasmids purchased from Addgene shown on table 5, virus generated by NU SBDRC). Cells were incubated with the virus on Matrigel-coated plates for 24 hours before removal of the virus. A subset of cells was maintained as virally transduced stem cells for expansion while the rest underwent differentiation.

### iPSC differentiation into Astrocytes (iA)

IPSC with rtTA, pTetO-Sox9-puro, and pTetO-Nfib-hygro lentiviruses were differentiated into astrocytes using a modified induction protocol previously published[54]. In short, cells were incubated in expansion media (DMEM, Thermo Fisher Scientific 11320082), 1x Glutamax (Thermofisher Scientific 35050-061), 1% N-2 supplement (Invitrogen, 17502-048), and 10% fetal bovine serum (FBS, Fisher Scientific, SH3007003) with 3ug/ml Doxycycline (Sigma, D3072) for 48 hours for lentiviral expression. Cells were gradually switched over 4 days into FGF media (Neurobasal media (Thermo Scientific, 12348017) containing 1X Glutamax 1X MEM nonessential amino acids (Life Technologies, 11140-050), 2% B27 supplement (Thermo Fisher Scientific, 17504044), and 1% FBS supplemented with 8ng/ml fibroblast growth factor (Peprotech, 100-18B), 10ng/ml BMP4 (R&D Systems, 314-BP), and 5ng/ml CNTF (Peprotech, 450-13) along with 1.25ug/ml puromycin (Sigma, P9620) and 200mg/ml hygromycin (Mirus Bio, MIR5930) for selection of lentiviral incorporated cells. Cells were then incubated in FGF media alone for 48 hours. On day 8, cells were replated at 250k cells/well on Matrigel-coated 12-well plates unless otherwise indicated and switched into astrocyte maturation media, 1:1 DMEM and Neurobasal media, 1% N-2 supplement, 1% Glutamax, 1% sodium pyruvate (Gibco, 11360070) supplemented with 5ng/ml heparin-binding EGF-like growth factor (Sigma-Aldrich, E4643), 0.1% Bovine serum albumin (Sigma-Aldrich, A7906), and 5mg/ml N-acetyl cysteine (Sigma-Aldrich, A8199). Cells were maintained with half-medium changes every 2 to 3 days until day 35 followed by treatment and collection for relevant analysis as described below.

### iPSC differentiation into Neurons (iN)

iPSC with lentiviral transduction of rtTA and pTetO-Ngn2-puro underwent differentiation via a well-described protocol for excitatory forebrain neurons with minor modifications[90]. The day following single-cell plating, cells were incubated in KO-DMEM media (Thermo Fisher Scientific, 10829018), 1X Glutamax, 1X MEM nonessential amino acids, and 0.1% 2-mercaptoethanol (Gibco, 21985023) that was supplemented with 10 uM SB431542 (Stemgent, 04-0010-05), 100 nM LDN193189 (Stemgent, 04-0074-02), 2 uM XAV939 (Tocris Bioscience, 3748), and 3 μg/ml doxycycline to induce lentiviral expression and subsequent NGN2 overexpression. Cells were then switched to 1:1 KO-media and neural induction media (NIM) containing 1x MEM nonessential amino acids, 1x Glutamax, 1x N-2, D-glucose (Sigma-Aldrich, G8769), 2ug/ml heparin sulfate (Sigma, H3149) supplemented with 3ug/ml doxycycline and 2ug/ml puromycin (Sigma, P9620). Following 24h incubation, the cells were switched into only NIM for 48 hours. Neural progenitor cells were resuspended using accutase and replated at 250k cells/well on Matrigel-coated 12-well plates unless otherwise indicated and maintained in BrainPhys neuronal media (Stemcell technologies, 05791) containing 1x MEM nonessential amino acids, 1x glutamax, 2% B27 supplement, 1% N-2 supplement, 10ng/ml BDNF (Peprotech, AF-450-02), and 3μg/ml doxycycline. The following day, cells were treated with 3uM Ara-C (Thermofisher Scientific, 449561000) for 48 hours to eliminate proliferating cells. iN were maintained with half-media changes every 2 to 3 days for 35 days until treatment and collection.

### iN and iA Coculture experiments

iA and iN were generated according to the protocols above. For direct culture experiments for single-cell RNA sequencing, iN and iA were plated 1:1 in 6-well Matrigel-coated plates for a total of 500k cells/well and maintained in 1:1 BrainPhys and astrocyte maturation media as described above until day 35. For indirect coculture experiments, neurons were plated at 250k cells/well in Matrigel-coated 12-well plates. Astrocytes were plated at 250k cells/well in Matrigel-coated Costar 12mm transwell inserts (Millipore Sigma, CLS3460) and maintained in 1:1 neuronal and astrocyte maturation media as described until day 35.

### iA and iN Treatments

Treatment of all differentiated cells was performed on day 35. All media was removed and fresh media with relevant treatment was added to the cells. For BMP4 treatments, cells were treated with 10ng/ml BMP4 (R&D Systems, 314-BP) for 72 hours before collection for analysis. The dosage and timing of BMP4 treatment were selected based on prior studies that determined the minimum required concentration for consistent SMAD phosphorylation in astrocytes and the duration necessary for reliable detection of BMP4 induced senescence marker expression. For autophagic inhibition, cells were treated with 200ng/ml Bafilomycin A1 (EMD Millipore, 196000) for two hours before collection. For Oxidized LDL experiments, 500ng/ml Oxidized low-density lipoproteins from human plasma (Thermo Scientific, L34357) were incubated on iN for 72 hours unless otherwise indicated. Astrocyte-conditioned media (ACM) was collected from iA following 72 hours of incubation and spun to collect cell debris before neuronal treatment. Lipid uptake inhibition experiments were performed concurrently with ACM treatments with 10μM lipid uptake inhibitor (Sigma-Aldrich, 5387690001).

### Single-cell RNA sequencing (scRNA-seq) Analysis

Cultured astrocytes were dissociated with accutase, pelleted at 0.8 RPM for 5 minutes, and resuspended in 0.04% BSA in PBS, followed by cell count. Cells were pelleted again at 300 RCF and resuspended in 100 μL of oligo multiplex tag, mechanically titrated via pipette 15 times, and incubated for 5 minutes, followed by the addition of 1.9 mL cold 1% BSA in PBS. Cells were spun at 300 RCF for 5 minutes at 4C. Supernatant was removed, and cells were resuspended and washed with 2 mL 1% BSA in PBS. Pellet and wash steps were repeated two more times. At final resuspension, cells were diluted in 1% BSA in PBS for a final concentration of 1,500 cells per μL based on cell count. Dissociated single cells were verified for greater than 80% viability and submitted for single-cell RNA sequencing using the Chromium 3’ v3.1 NexGem Single Cell platform (10x Genomics) with assistance from the Northwestern University genomics core. 30,000 cells from three to six independent wells per experimental condition were combined, captured and barcoded on a microfluidic chip, subsequently used for cDNA library construction before sequencing on the Illumina NovaSeq 6000 S4 platform with 150bp pair-end reads. The FASTQ formatted sequencing data were pre-processed using Cell Ranger v7 or Cloud Analysis (10x Genomics) for each sample and aligned with the GRCh38 human reference genome and the number of reads from each cell that align to each gene were quantified. The matrix files which summarize the alignment results and were then imported in Seurat (v5, Satija Lab, NYGC) for further analysis. QC of each experimental sample was independently performed to exclude cells with fewer than 200 or greater than 3000 genes and greater than 10% mitochondrial DNA content to poor-quality cells. Data from all samples underwent log normalization and scaling followed by variable feature analyses before and after experimental sample integration via the Harmony R package [91]. Top 10 PCs of the Principal component analysis results were used for subsequent nearest neighbor and cluster identification using snn resolution of 0.2. Cell cluster identities were validated by expression of known cell lineage-specific markers, with finalized distribution visualized in two dimensions using uniform manifold approximation and projection (UMAP) dimensional reduction. Identification of cell cluster and experimental group-specific differentially expressed genes (DEG) were performed using the Seurat functions *FindMarkers* or *FindAllMarkers* with fold change threshold of 1.5 and min.pct value of 0.25. Both up and down-regulated DEGs were used for pathway analyses performed via Enrichr[56] and confirmed by GSEA analysis using the clusterProfiler R package[57]. Expression of genes of interest on the UMAP was presented as using the *FeaturePlot* function in Seurat. The Seurat function *AggregateExpression* was used to perform pseudobulk expression quantification for each experimental condition, which are used for relative gene expression comparisons across experimental group of specific genesets of interest. *AddModuleScore* function in Seurat was used to assess normalized expression of specific genesets of interest across experimental groups. Cell-cell communication inference was performed using CellChat R package[92].

### RNA extraction and Quantitative Real-time PCR (qPCR)

Cells were single cell dissociated with accutase for 15 minutes followed by 1:1 quenching with appropriate astrocyte or neuronal media. Cells were collected into a 15ml falcon conical tube (Corning, 352070) and spun at 0.8 RPM for 5 minutes. The supernatant was discarded, and cells were resuspended with cold 1xDPBS (Corning, 21-031-CV). RNA extraction was conducted using the RNAqueous Micro Total RNA Isolation Kit according to the manufacturer’s instructions (Thermo Fisher Scientific, AM1931) and eluted in 25uL ultra-pure H20. Quality control and quantification of RNA eluted was done using a Nanodrop 2000c spectrophotometer. Reverse transcription of resultant RNA was performed according to kit protocol using the SuperScript IV VILO Master Mix (Thermo Fisher Scientific, 11766050). SYBR Green PCR Master Mix (Thermo Fisher Scientific, 4309155) and MicroAmp Optical 384-well reaction plates (Thermo Fisher Scientific, 4309849) were used to perform real-time quantitative PCR (qPCR). The primers used for ID3 were as follows: Forward, 5′-GAGAGGCACTCAGCTTAGCC -3′; Reverse, 5′- TCCTTTTGTCGTTGGAGATGAC - 3′.The NUSeq core provided the QuantStudio 7 Flex Real-time PCR system for qPCR. Assessment of threshold cycle (Ct) for target genes and GAPDH as the housekeeping gene were used to calculate the relative expression of each gene (ΔCt). ΔCt of all groups were compared to ΔCt of E3 vehicle-treated iAs to determine ΔΔCt values and were expressed as fold changes.

### Protein analysis

#### Protein isolation and quantification

Cultured cells were washed with room temperature (RT) 1x DPBS (Corning, 21-031-CV) followed by Pierce m-Per protein extraction reagent (Thermo Scientific, 78505) supplemented with 1mM Halt protease and phosphatase inhibitor cocktail (Fisher Scientific, 78444) before removal of cells from the plate using a cell scraper (Fisher Scientific, 08-771-1A). Cells were broken up via mechanical trituration and transferred to a sterile microcentrifuge tube (VWR, 10011-724) and placed on ice. Protein was vortexed for 30 seconds every five minutes for a total of 30 minutes. Cells were then spun at 14,000 RPM at 4C for 20 minutes to pellet cellular debris. Protein quantification of the supernatant was performed via Pierce Rapid Gold BCA (Thermo Fisher Scientific, A53226) kit instructions utilizing a Biotek Synergy HTX microplate reader.

#### Immunoblotting

Cellular protein was run on 4-20% tris-glycine gels (Bio-Rad Laboratories, 4561085) and transferred onto Immobilon polyvinylidene difluoride membranes (PVDF; Millipore, IPVH00010). The membranes were blocked in 5% BSA for one hour. Primary antibodies were incubated overnight at 4C on an orbital shaker (Table 1 includes a list of antibodies used). After primary incubation, membranes were washed three times with 1x tris buffered saline (Bio-Rad, 1706435) before incubation in secondary antibody for one hour at RT on an orbital shaker. Secondary antibodies used were horse anti-mouse horseradish peroxidase (Cell Signaling Technology, 7076) and goat anti-rabbit horseradish peroxidase (Cell Signaling Technology, 7074). For chemiluminescent imaging, blots were briefly incubated in Radiance Q substrate (Azure Biosystems, AC2101) and developed using the Azure c600 bioanalytic imaging system. Quantification of blots was performed using ImageJ software (NIH, Bethesda, MD, USA). Protein expression was evaluated in triplicate across 3 independent differentiations.

**Table 1.** List of astrocytic markers used to determine the maturity score of iAs subclusters in scRNAseq.

### Immunocytochemistry and Confocal imaging

Cell cultures were differentiated as previously described and plated at a density of 100k per well on Matrigel-coated coverslips (Azure Scientific, ES0117520) and treated with BMP at day 35 for 72 hours. For fixation, cells were washed once with 1x dPBS before being fixed with 4% Paraformaldehyde for 20 minutes. Coverslips were subsequently washed three times with 1x dPBS. For immunocytochemical staining, cells were blocked in 5% normal goat serum (Jackson ImmunoResearch, 005-000-121), 1% BSA, 0.25% Triton X-100 (Sigma-Aldrich, T8787) in PBS for 1 hour at RT. Primary antibodies, found in Table 2, were diluted in blocking solution and incubated overnight at 4C. The following day, coverslips were washed three times with 1x PBS, 0.25% Triton-X100 (PBST) before incubation in secondary antibody mix, LipidTOX neutral lipid stain (Thermo Scientific, H34476), and Hoescht 33342 (Thermo Fisher Scientific, 62249) for one hour at RT. Coverslips were washed five times in 1X PBST and mounted with Prolong Diamond Antifade Reagent (Thermo Scientific, 3970) onto Superfrost Plus microscope slides (Fisher Scientific, 12-550-15). Confocal imaging was done using a Leica SP5 Confocal microscope with Z-stacks (step size= 1 μm). 5 or more images were quantified for each experiment per condition. Analysis of lipid droplet colocalization or size was performed using Cell Profiler software[93]. Analyses were conducted across 5 independent experiments with at least 5 images acquired per experiment.

**Table 2.** List of Antibodies used.

| **Antibodies** | **Source** | **Catalog** |
| --- | --- | --- |
| S100B anti-ms | Sigma-Aldrich | S2532 |
| GFAP anti-rb | DAKO | Z033401-2 |
| GLT-1 | Santa Cruz | sc-365634 |
| Connexin-43 (F-7); anti-ms | Santa Cruz | sc-271837 |
| Vimentin; anti-ck | Millipore | AB5733 |
| p-SMAD1/5/8; anti-rb | Millipore | AB3848-1 |
| t-SMAD | ABCAM | AB13723 |
| GAPDH (G-9); anti-ms | Santa Cruz | Sc-256062 |
| LipidTOX | Thermo Fisher Scientific | H34476 |
| DAPI | Thermo Fisher Scientific | 62249 |
| LAMP1; anti-rb | Sigma-Aldrich | L1418 |
| LAL; anti-ms | ABCAM | AB89771 |
| ATGL; anti-rb | Cell Signaling Technology | 2138S |
| LC3; anti-rb | Sigma-Aldrich | L8918 |
| Amyloid; anti-ms | Biolegend | SIG-39300 |
| TUJ1; anti-rb | ABCAM | AB52623 |
| AT8; anti-ms | Thermo Fisher Scientific | MN1020 |
| HT7; anti-ms | Thermo Fisher Scientific | MN1000 |
| pP38; anti-rb | Cell Signaling Technology | 9211S |
| P38; anti-rb | Cell Signaling Technology | 9212S |

**Table 3.**
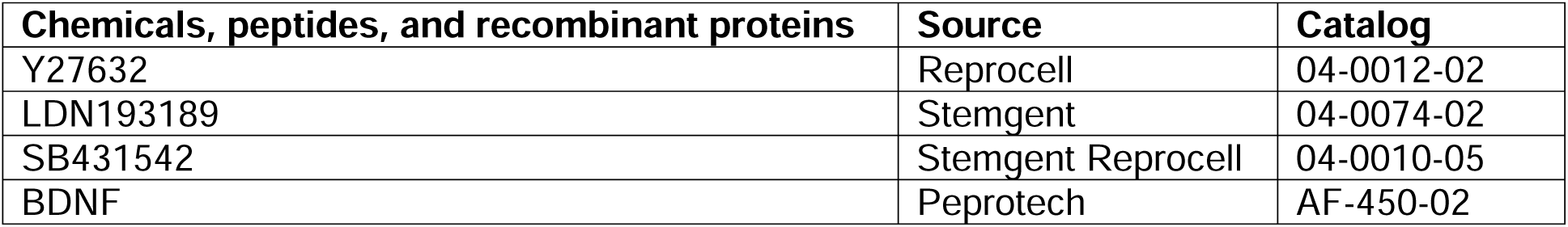
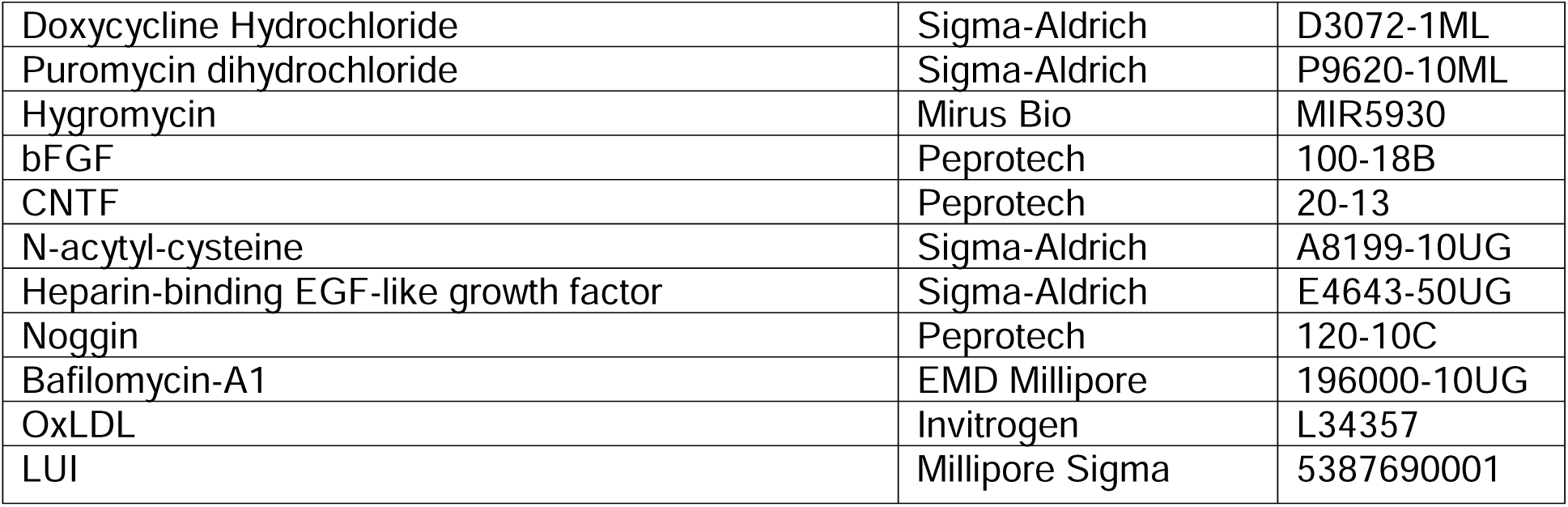
List of chemicals, peptides, and recombinant proteins.

**Table 4.**
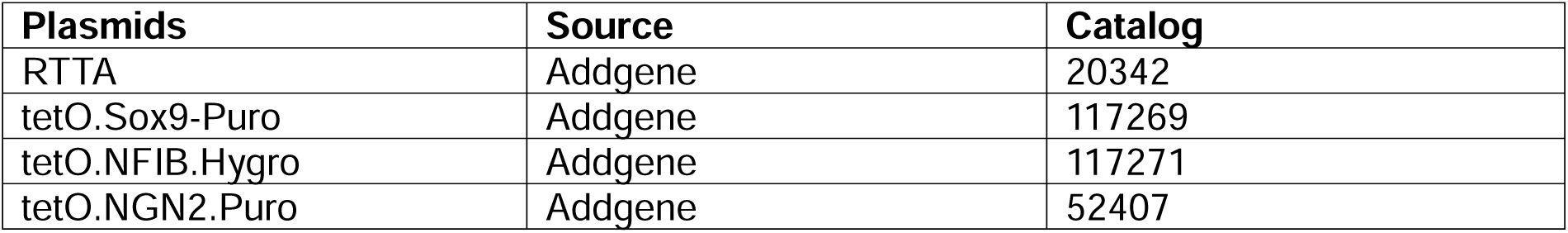
List of plasmids.

**Table 5.** List of iPSC cell lines.

| Human iPSC cell lines | Source |
| --- | --- |
| Human induced pluripotent Stemcell<br>Line: AG011414 | Wadhwani et al |
| Human induced pluripotent Stemcell<br>Line: Clone G10 | Wadhwani et al |
| Human induced pluripotent Stemcell<br>Line: AG05810 | Wadhwani et al |
| Human induced pluripotent Stemcell<br>Line: Clone D6 | Wadhwani et al |
| Human induced pluripotent Stemcell<br>Line: AG04402 | Affaneh et al |
| Human induced pluripotent Stemcell<br>Line: Clone H7 | Affaneh et al |

### Lipid isolation

Treated cells were washed with 1x PBS followed by accutase dissociation as described above and transferred to a 1.5mL microcentrifuge tube. Supernatant was removed and cells were resuspended in PBS followed by cell pelleting at 1000g for 5 minutes. Cells were resuspended in 25 μL of PBS and lipid extraction was performed according to protocol of lipid extraction kit (Abcam, ab211044). 500 μL of lipid extraction buffer was added to the cells and tubes were agitated on an orbital shaker at room temperature for 20 minutes. The tubes were then spun at 10,000g for 5 minutes and supernatant was transferred to a new tube. The supernatant was then dried out in a 37C degree incubator overnight and contents in the tube were resuspended in suspension buffer before being analyzed for triglyceride or cholesterol components.

### Cholesterol assay

Samples were assessed for cholesterol components according to HDL and LDL/vLDL Cholesterol Assay Kit instructions (Cell Biolabs, STA-391). In short, HDL and LDL fractions were split by adding precipitation reagent to each sample and vortexing, followed by a 10-minute incubation at room temperature. Samples were spun at 2000g for 20 minutes and the HDL-containing supernatant was transferred to a new tube. The pelleted LDL fraction was dissolved in PBS and vortexed to dissolve. Samples and standards were added to a 96-well microtiter plate with 1:1 cholesterol reaction reagent with and without cholesterol esterase for detection of total, esterified, and free cholesterol. Contents were mixed followed by incubating the plate at 37C for 45 minutes without light. Fluorescence was read using a Biotek Synergy HTX at excitation of 530 and emission at 590. Standard curves were used to calculate levels of HDL and LDL cholesterol species within the sample. Samples were performed in duplicate and averaged across conditions. Analyses was performed in triplicate across three independent experiments.

### Triglyceride assay

Cells and media were assayed according to Triglyceride Glo kit instructions (Promega, J3160. Glycerol lysis solution with and without lipase was added to samples and shaken before incubated at 37C for 30 minutes and transferred to a 96-well plate along with standard dilutions. Glycerol detection reagents were added to the plate and shaken briefly followed by incubation at RT for 1 hour. Luminescence of standards and samples were determined using a Biotek Synergy HTX and quantified levels of triglycerides based on the difference between free and total glycerol levels in the sample. Samples were performed in duplicate across three independent experiments and averaged across conditions. Subsequent statistical analysis of values was performed as described below.

### Enzyme-linked Immunosorbent Assay (ELISA)

72-hour treated conditioned iA media was collected from 12-well plates and treated with Halt Protease and phosphatase inhibitor followed by centrifugation to collect cellular debris. For cellular levels of target protein, cells were collected and prepared with lipid extraction or protein extraction as described above. The samples were analyzed via Human ApoE Elisa Kit (Cell BioLabs, STA-367) or OxiSelect Human Oxidized LDL Elisa Kit (Cell BioLabs, STA389). Sample preparation and assay protocol was performed according to kit instructions and colorimetric quantitation was performed using Biotek Synergy HTX at 450nm. Samples were performed in duplicate across three independent experiments. Subsequent statistical analysis of values was performed as described below.

### Cytokine Array

Detection of cytokine expression in ACM was conducted using Proteome Profiler Human Cytokine Array (R&D Systems, ARY005B). ACM was collected following 72-hour Vehicle or BMP treatment, treated with Halt Protease and Phosphatase inhibitor, and centrifuged to collect cellular debris. Cytokine array was conducted according to kit instructions. Chemiluminescent signal of membranes was visualized using Azure c600 bioanalytic imaging system exposure for 10 minutes and relative change of cytokine expression was quantified using ImageJ software. Subsequent statistical analysis was performed as described below.

### Seahorse analysis

iA were plated on 96 well plates (Fisher Scientific, 08-772-2C) at a density of 30,000 cells per well, with corner wells omitted for background reference. Following 72 hours of vehicle or BMP treatment, Seahorse analysis was conducted using Seahorse XF Cell Mito Stress Test Kit (Agilent, 103015) or XF Palmitate Oxidation Stress Test Kit (Agilent, 103693-100) according to kit protocols. For fatty acid oxidation kit, cells were transitioned one day prior to assay into substrate-limited growth media containing 0.5mM glucose, 1mM glutamine, 1% FBS, 0.5 mM L-Carnitine with relevant treatment conditions preserved and incubated overnight in 37C incubator with 5% CO2. Calibration and hydration of sensor cartridges were performed according to kit instructions. For the day of assay, cells were washed according to kit protocols, with relevant treatment still included. Cells were incubated in assay diluent based on kit protocols in a 37C non-CO2 incubator for one hour. Fresh assay media was added immediately before plate was loaded into a XFe96 Seahorse Analyzer available through Northwestern University Metabolomics Core Facility (NU-MCF). Port loading in the sensor cartridges was performed according to kit instructions, with 10X final concentrations of Etomoxir, Oligomycin, FCCP, or Rotenone/Antimycin A depending on kit instructions. Final concentrations of drugs treated for iA seahorse were: 4μM etomoxir, 1.5μM oligomycin, 2μM FCCP, and 0.5μM Rotenone/Antimycin A. Additional analysis was performed using Seahorse Wave Pro Software and online software client (https://seahorseanalytics.agilent.com/).

### IncuCyte imaging

iN cultures were treated with Vehicle, BMP4, oxLDL, or LUI or oxLDL + LUI at concentrations listed above. Live and dead cell analysis was conducted using 10nM Calcien AM (Fisher Scientific, C3099) and 250nM of Cytotox Red (Essen Bioscience, 4632). Treated cell plates were loaded into a Sartorius IncuCyte S3 Live Cell Imaging system (Essen Bioscience) available through the Northwestern University Analytical BioNanoTechnology Equipment Core (NU-ANTEC). Brightfield and Green (excitation 440-480, emission 504-544)/red (excitation 565-607, emission 625-705) fluorescence was imaged at 20x (0.62 μm/pixel) every 6 hours for 72 hours with 9 fields per well. Images had a 2% spectral unmixing of red from green channel and quantification of live and dead cell number was conducted using Sartorius IncuCyte analysis software. Live cell over total cell counts were averaged per well at each time point and normalized to time 0h. Quantification of live cells per time point was expressed as a percent of total cells in the image frame across fields. Subsequent analysis was performed via two-way repeated measures ANOVA with Tukey test for multiple comparisons.

### Statistical analysis

Data collected were conducted across independent patient lines N=3 wells per experiment. Data analysis was conducted using Prism 10 software (GraphPad) and all results are reported as mean ± SEM. Statistical analysis was conducted for groups using two-way analysis of variance (ANOVA) followed by Tukey’s post hoc test under normal distribution. For time-dependent Seahorse and IncuCyte experiments, statistical analysis was conducted via two-way ANOVA with repeated measures followed by Tukey post hoc test for multiple comparisons at each time point. Statistical significance was predetermined with an assigned cut-off at p<0.05.

## Supplemental Information titles and legends

**Supplemental Figure 1: iA differentiation generated mature astrocyte populations that are responsive to BMP4 treatment.** A) Schematic of lentiviral constructs used for iA generation B. Representative confocal images of relevant mature astrocytic markers stained via immunocytochemistry: S100B, GFAP, GLT-1, Connexin-43, and Vimentin (Scale bar = 100μm). C) Quantification of glutamate uptake of iA showing no difference across groups (n=6). D) Relative qPCR transcript levels of ID3 following vehicle or BMP treatment in iA cultures (n=3). E) Lipid droplets per cell binned by size up to 3μm across all culture conditions (n=5). F) Representative western blot and quantification of APOE protein (n=8). G) Quantification of APOE secreted into astrocyte-conditioned media based on ELISA (n=9). Data are expressed as mean ± SEM using two-way ANOVA followed by Tukey post-hoc test for multiple comparisons to calculate significance (*p<0.05, **p<0.01, ***p<0.001, ****p<0.0001).

**Supplemental Figure 2. Analysis of induced-astrocyte single-cell RNA sequencing results confirmed astrocytic identity and enrichment in cholesterol biosynthesis in E4 BMP iA.** A) UMAP representation of cell cluster distribution from the iA scRNAseq results with the expression profiles of select cell state and lineage-specific molecular markers including SOX9, NFIA, HOPX for glial precursors, GFAP, S100B, CD44, APOE, ITGA6 for astrocytes, MKI67 for progenitors in cell cycle, DCX and SYN1 for neuronal lineage-committed cells. B) Heatmap representation of relative expression of pan-reactive, A1-reative, and A2-reactive astrocyte markers across the 4 experimental groups illustrating high A2-reactive astrocyte marker expression in cluster Ast4 cells. C) Venn diagram illustrating the number of total, unique, and shared differentially down-regulated genes in the 4 experimental groups. D-F) Differentially regulated pathways identified in pairwise comparisons using Gene Set Enrichment Analysis (GSEA) between (D) E3 Veh and E3 BMP (E), E3 BMP and E4 BMP, and (F) E4 Veh and E4 BMP experimental groups. No enriched pathways were detected in E3 Veh and E4 Veh. (G) Heatmap presentation of relative expression of fatty acid biosynthesis genes across the 4 experimental groups. (H) Violin plot representation of normalized expression of fatty acid biosynthesis genes compared across the 4 experimental groups. I) APOE mRNA expression by scRNAseq (I) and protein expression by western blot (J) in the 4 experimental groups.

**Supplemental Figure 3: Cholesterol synthesis and metabolism genes are upregulated in E4 iA.** A) Quantified levels of total, HDL, and vLDL/LDL cholesterol within iA cells in uM (n=6). B) Representative western blot and quantification of FASN protein expression in iA (n=6). C) Representative western blot and quantification of SREBP2 protein in iA (n=6). D) Representative western blot and quantification of LCAT protein in iA (n=6). E) Representative western blot image and quantification of CETP in iA (n=6). Data are expressed as mean ± SEM using two-way ANOVA followed by Tukey post-hoc test for multiple comparisons to calculate significance (*p<0.05, **p<0.01, ***p<0.001, ****p<0.0001).

**Supplemental Figure 4: Mitochondrial respiration is modulated by BMP signaling in iA cultures.** A) Extracellular acidification rate (ECAR) of iA cultures during seahorse assay in mpH/min (n=8). B) Average ECAR of iA cultures across seahorse assay timeline shown in A. C) Basal ATP production rate through both mitochondrial and glycolytic pathways based on seahorse analysis. Blue bars: MitoATP; Red bars: GlycoATP (n=8). D) Baseline etomoxir response in iAs cultures between vehicle and etomoxir treated groups in pmol/min (n=6). Data are expressed as mean ± SEM using two-way ANOVA followed by Tukey post-hoc test for multiple comparisons to calculate significance (*p<0.05, **p<0.01, ***p<0.001, ****p<0.0001).

**Supplemental Figure 5. BMP signaling does not significantly alter the expression of astrocyte-secreted cytokines and b-amyloid proteins.** A-E) Immunoblot densitometry quantification of select astrocyte-expressing cytokines including (A) CCL2, (B) MIF, (C) IL6, (D)IL8, (E) CXCL1, detected in the ACM from the E3/E4 iA cultures treated with either vehicle or BMP4 (n=6). No significant changes were observed between BMP treatment conditions in iA with the same APOE genotype. (F, G) Representative western blot and quantification of beta-Amyloid protein (n=3). (H-J) ELISA of amyloid beta protein isoforms in ACM showing BMP signaling does not differentially regulate cellular and secreted Ab levels (n=9). Data are expressed as mean ± SEM using two-way ANOVA followed by Tukey post-hoc test for multiple comparisons to calculate significance (*p<0.05, **p<0.01, ***p<0.001, ****p<0.0001).

**Supplemental Figure 6. Single-cell RNA sequencing of induced-neurons revealed BMP signaling reduces expression of oxidative phosphorylation-related transcripts.** A) UMAP representation of cell cluster distribution from the iN scRNAseq results with the expression profiles of select cortical excitatory neuron markers including MAP2, SLC17A6 (vGlut2), LRP1B, PHOX2B, GRM7, GRIA2, RORB, and ISL1. B) Top 10 enriched pathways identified using the Wikipathway2024 genesets in vehicle C) Violin plot representation of normalized expression of 13 known BMP target genes confirming the efficacy of BMP4 treatment in iNs. D) Violin plot representation of normalized express of 33 known cholesterol biosynthesis genes showning no differential expression between experimental groups. ***p<0.001.

**Supplemental Figure 7. Single-cell RNA sequencing of induced astrocyte-neuron cocultures revealed paracrine signaling across cell types.** A) UMAP representation of cell cluster distribution from the iN+iA coculture scRNAseq results with the expression profiles of select cell type-specific markers including neural precursor markers MKI67, HOPX, SOX9, astrocyte markers GFAP, S100B, CD44, APOE, and neuronal markers SLC17A6, GRIA2, ISL1, TBR1. B) Top 5 enriched pathways identified using the Wikipathway2024 genesets in the iA of the 4 coculture experimental conditions. Red box highlights the enrichment of transcripts related to cholesterol synthesis in the cocultured E4 BMP iA. C) Top 5 enriched pathways in E3iN cocultured with E3iA in the presence and absence of BMP4 using the Wikipathway2024 genesets.

Supplemental Table 1. List of enriched DEGs and pathways in BMP4 or vehicle treated E3 and E4 induced astrocytes.

Supplemental Table 2. List of enriched DEGs and pathways in BMP4 or vehicle trated E3 induced neurons.

Supplemental Table 3. List of enriched DEGs and pathways in BMP4 or vehicle treated E3 or E4 induced astrocytes from iN+iA coculture experimental gorups.

Supplemental Table 4. List of enriched DEGs and pathways in BMP4 or vehicle treated E3 induced neurons from 4 iN+iA coculture experimental groups.

