## Supplementary figures and images for "Differential Regulation of APOE4-Mediated Astrocytic Lipid Metabolism by BMP Signaling Exacerbates Alzheimer’s Disease Pathologies in Neurons"

### Figure S1

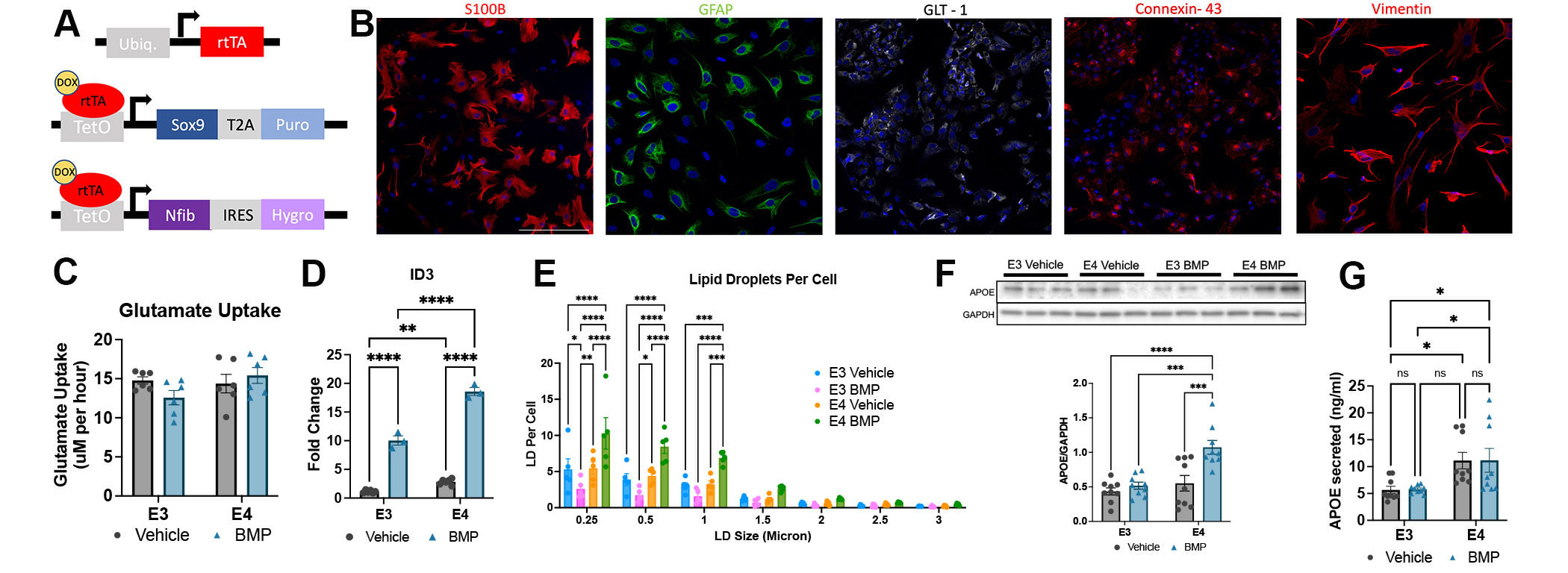

### Figure S2

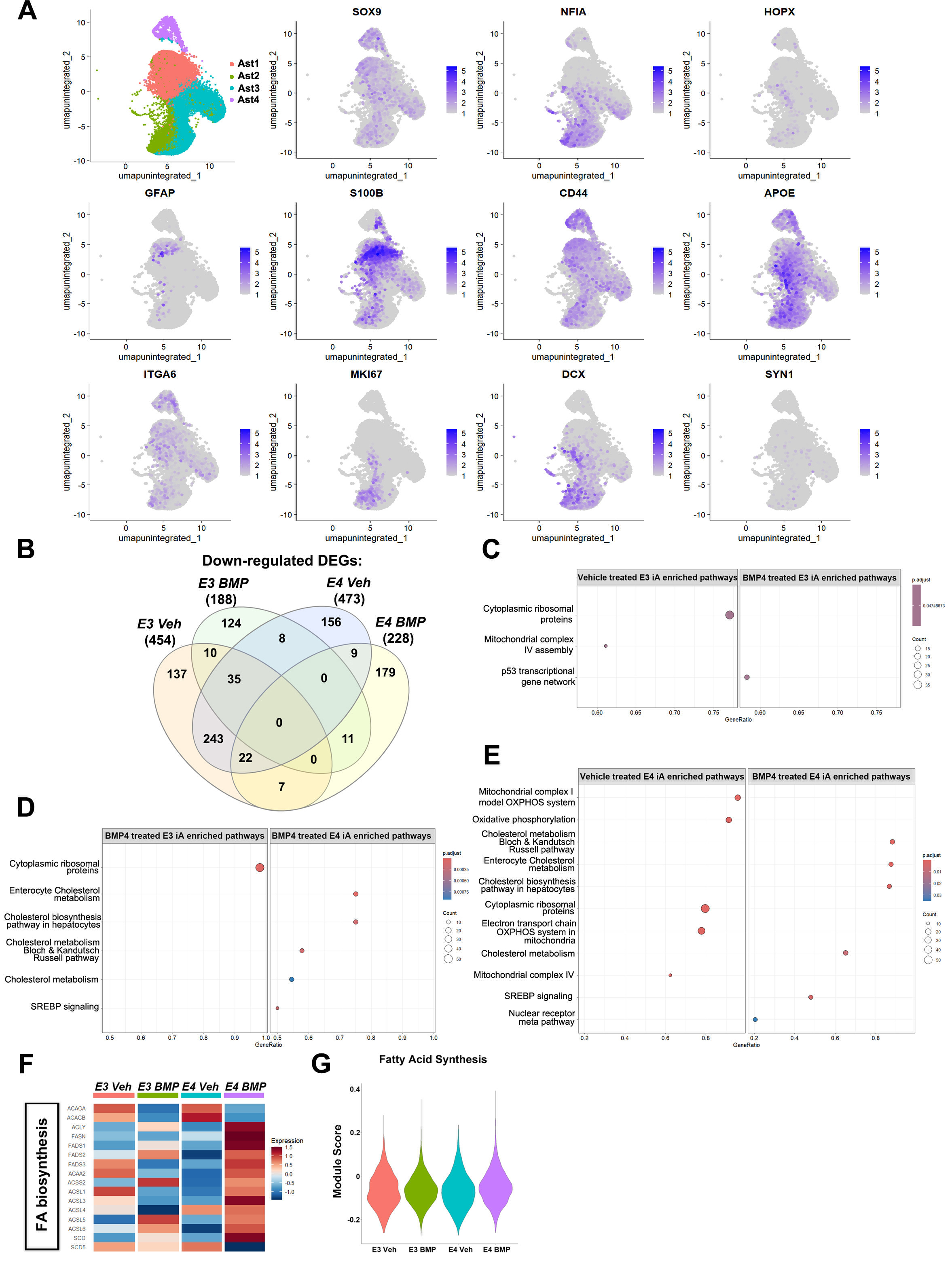

### Figure S3

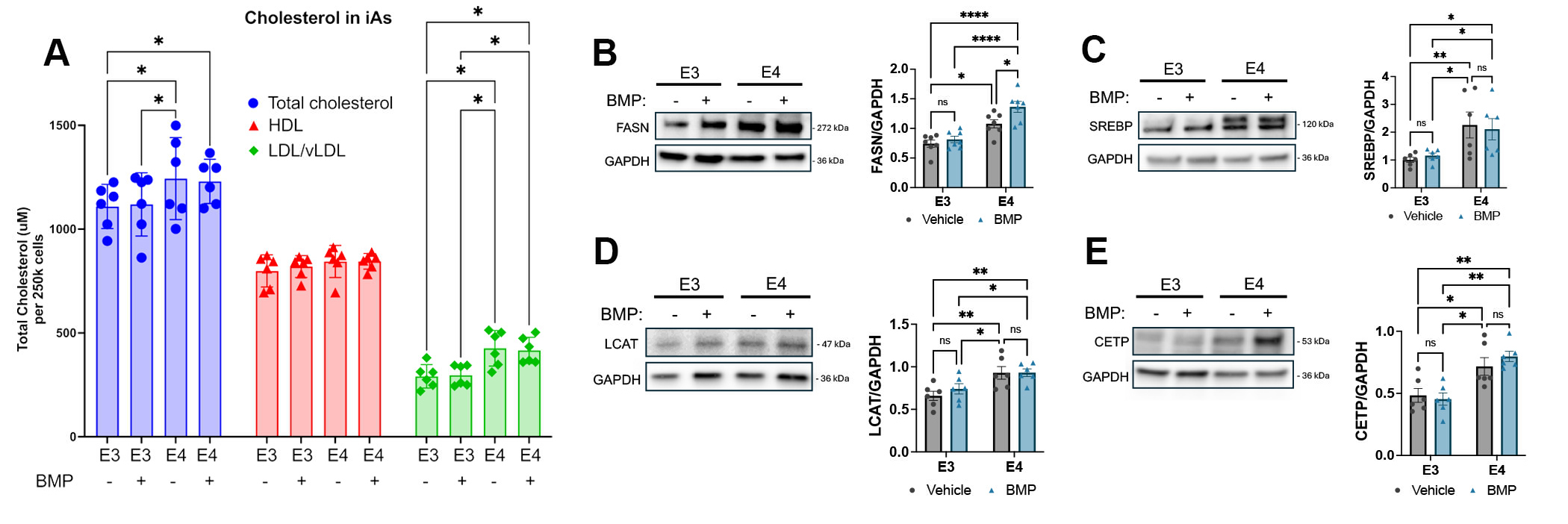

### Figure S4

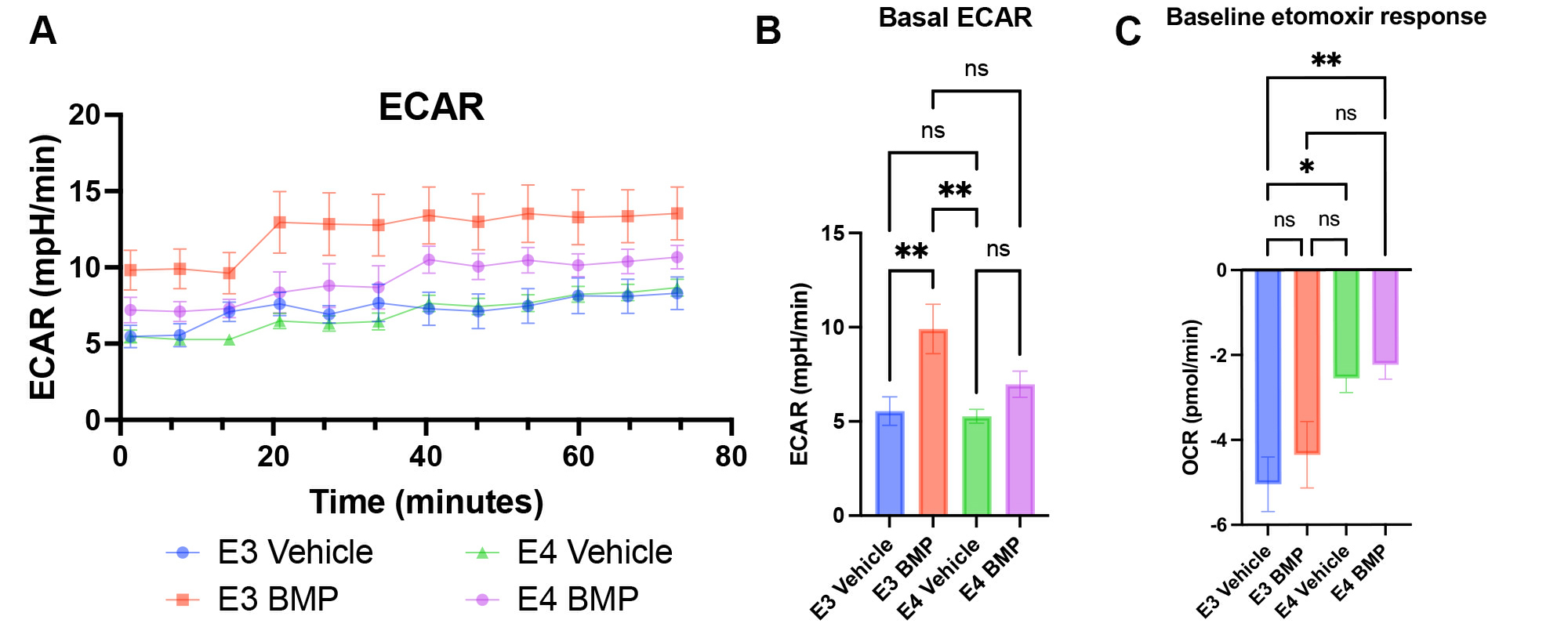

### Figure S5

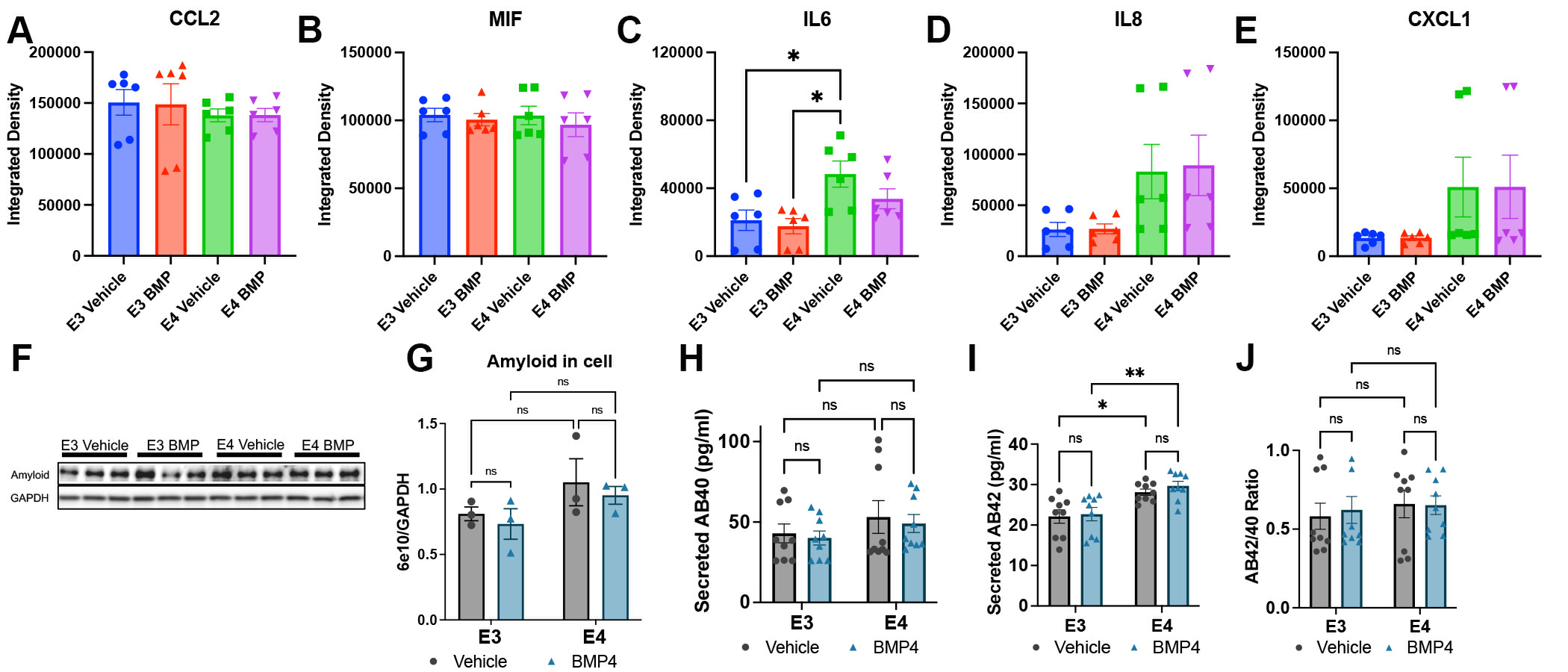

### Figure S6

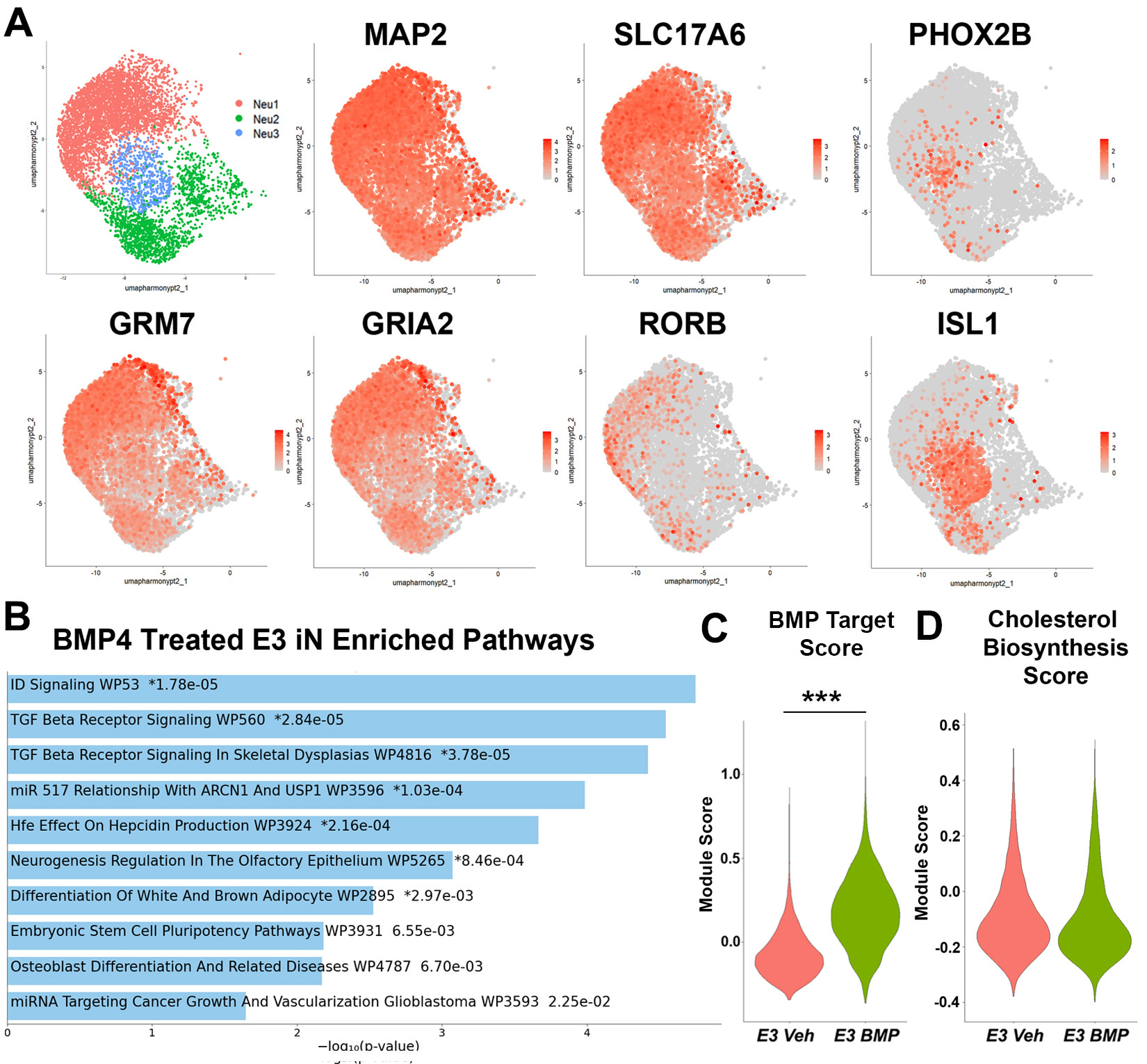

### Figure S7

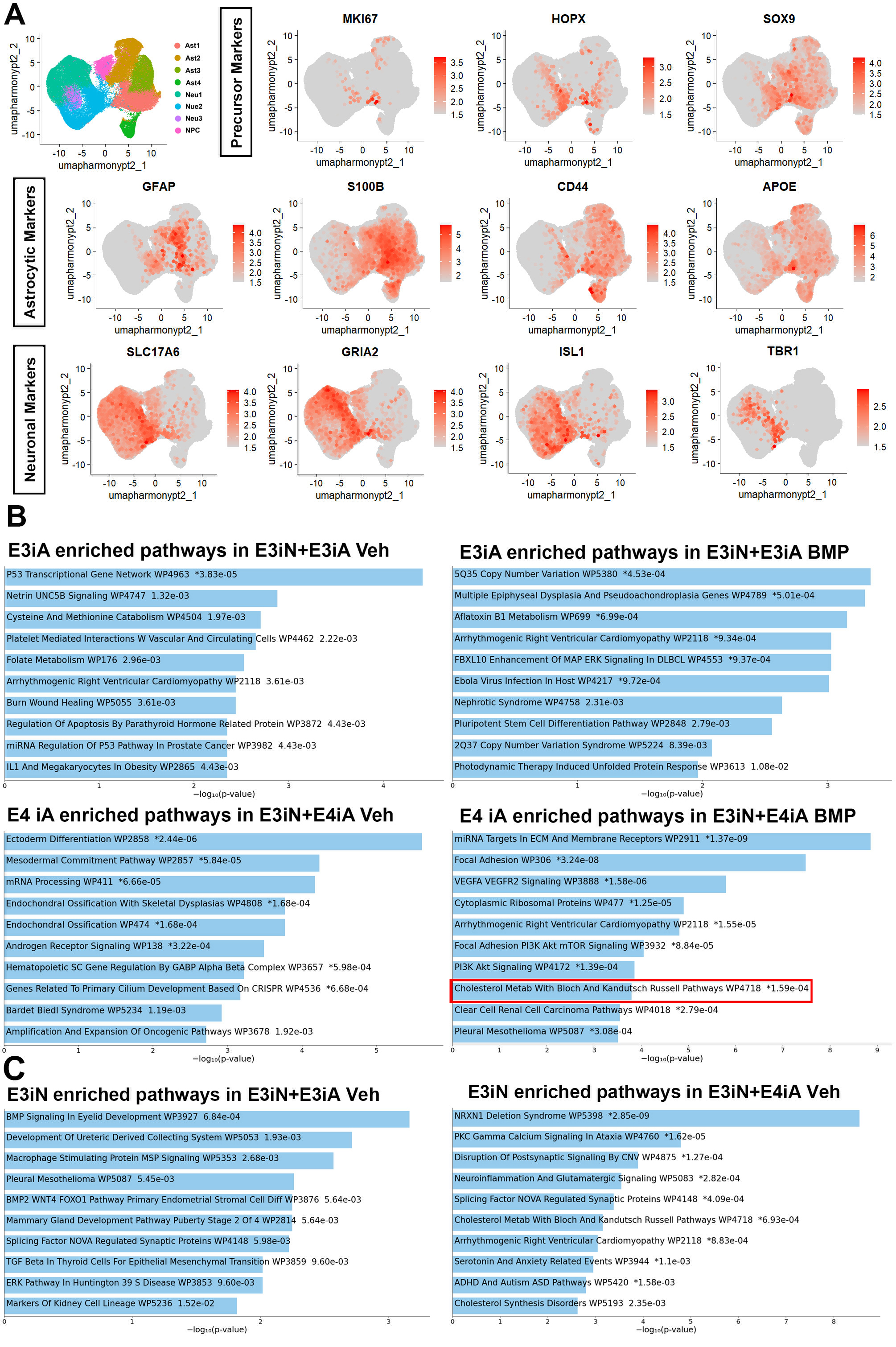
